# *In vivo* functional phenotypes from a computational epistatic model of evolution

**DOI:** 10.1101/2023.05.24.542176

**Authors:** Sophia Alvarez, Charisse M. Nartey, Nicholas Mercado, Alberto de la Paz, Tea Huseinbegovic, Faruck Morcos

**Author notes:** **Materials and Correspondence** Correspondence to Faruck Morcos at. These authors contributed equally to this work (Sophia Alvarez, Charisse M. Nartey).

## Abstract

Computational models of evolution are valuable for understanding the dynamics of sequence variation, to infer phylogenetic relationships or potential evolutionary pathways and for biomedical and industrial applications. Despite these benefits, few have validated their propensities to generate outputs with *in vivo* functionality, which would enhance their value as accurate and interpretable evolutionary algorithms. We demonstrate the power of epistasis inferred from natural protein families to evolve sequence variants in an algorithm we developed called Sequence Evolution with Epistatic Contributions. Utilizing the Hamiltonian of the joint probability of sequences in the family as fitness metric, we sampled and experimentally tested for *in vivo β*-lactamase activity in *E. coli* TEM-1 variants. These evolved proteins can have dozens of mutations dispersed across the structure while preserving sites essential for both catalysis and interactions. Remarkably, these variants retain family-like functionality while being more active than their WT predecessor. We found that depending on the inference method used to generate the epistatic constraints, different parameters simulate diverse selection strengths. Under weaker selection, local Hamiltonian fluctuations reliably predict relative changes to variant fitness, recapitulating neutral evolution. SEEC has the potential to explore the dynamics of neofunctionalization, characterize viral fitness landscapes and facilitate vaccine development.

## Introduction

Important features of protein structure, their functional capabilities, and the constraints imposed during the course of evolution can, in principle, be elucidated from sequence data and used to develop models of sequence evolution. The value of such models rests in the fact that these tools help us to understand not only past events, but the driving forces of protein-sequence change. Traditionally, models characterize subsets of statistical features found in natural sequence data, often requiring the application of multiple theories and practices to paint a comprehensive picture of evolution. We developed a model that unifies such features and uses them to guide unexplored evolutionary trajectories for sequences in specific protein families [1]. This model called Sequence Evolution with Epistatic Contributions (SEEC) utilizes a global inference model to recapitulate family sequence statistics determined from evolutionarily related epistatic interactions. The algorithm proceeds to sequentially evolve novel protein sequences with the potential to retain wild type (WT) functionality based on a conditional probability that takes into account epistasis and the sequence context at each evolutionary step. Beyond *de novo* protein design, our model provides insights on how one single protein can explore functional sequence space not yet discovered in natural data sets.

The SEEC model incorporates epistatic information provided by direct coupling analysis (DCA) [2], a joint probability covariance model that utilizes both pairwise and single-site statistics to infer the family couplings (*e*_*ij*_) and local fields (*h*_*i*_) parameters of the Potts model of the protein family sequence space [3, 2]. SEEC models neutral evolution by exploring new sequence space while preserving family-like function. In these simulations, SEEC unifies various evolutionary models with epistasis and the emergent properties of overdispersion, Gamma distribution of substitution rates across sites, heterotachous sites, and evolutionary Stokes shifts or entrenchment [1]. To demonstrate the significance of epistatic constraints, other groups have found that the application of coevolutionary information within molecular binding affinity highlighted the incorporation of epistasis and a changing mutational fitness which better modeled the dynamics of antibody evolution [4], while the development of an epistatic inference model that utilized time-series genetic data better simulated complex selection that can be applied to the analysis of virus, bacteria, and cancer allele evolution [5]. Entrenchment is another epistatic sequence evolution phenomenon that has been recently observed experimentally [6, 7]. Another key aspect of the SEEC evolutionary simulations involves the predictive power of the sequence Hamiltonian, a statistical energy calculated for each evolved sequence based on the probability of retained family-likeness. Many studies have explored the interconnection between the Hamiltonian, free energy landscapes, and protein fitness showcasing how family alignments encode biological information such as folding [8, 9], melting [10], mutational space [11], and molecular interaction potential [12]. Subsequently, the Hamiltonian energies are often correlated with distinguishing fitness such that sequences with lower statistical energies are more probable in regards to the landscape of the family [13]. Experimental data also shows strong correlations between empirically measured fitness differences and changes within the Hamiltonian energy of a given family [14, 15, 16]. The retention of these aforementioned evolutionary statistical features affirms the power of the Hamiltonian fitness landscape and its use in understanding the dynamics of sequence-function adaptation [17]. Altogether, the potential for SEEC to produce functional proteins motivates our rationale to further assess this evolutionary model with a direct experimental counterpart.

In order to test the capabilities of this model of epistatic evolution, we focused on the *E. coli* antibiotic resistance protein TEM-1 (UniProt ID P62593). The *β*-lactamase family is a convenient biological system for testing evolutionary models due to the ease of assaying protein activity with survival in the presence of antibiotics. Additionally, the plethora of sequence information and previous research available including the successful analysis of coevolutionary data for the *β*-lactamase family [15, 18, 19] makes this an ideal system to assess the performance of SEEC experimentally. We utilized SEEC to computationally evolve TEM-1 using parameters inferred from both mean-field and Boltzmann machine learning models [2, 20]. We identified and synthesized key variants that, when expressed from a plasmid, protected *E. coli* from ampicillin on par with or even better than the WT enzyme. Remarkably, some of these successful variants undergo about 448 substitutions and reversions leading to 47 point mutations when compared to the WT TEM-1 *β*-lactamase. The number of potential sequences created with the same magnitude of random mutations is enormous, and requires a trustworthy model to navigate this unexplored mutational space. Being able to predict functional sequences evolved dynamically from extant sequence data is a powerful tool; as such, SEEC exemplifies a model of neutral evolution capable of preserving the statistics of observed evolution as well as specifying generated sequences with the functional characteristics of its ancestors.

## Results

In order to assess SEEC’s evolutionary modeling experimentally, we first simulated computational protein evolution (Fig. 1a). To do this, we compiled an MSA for the antibiotic resistance family of *β*-lactamase enzymes (PF13354) which is used as the input for direct coupling analysis (DCA) to infer the family couplings (*e*_*ij*_) and local fields (*h*_*i*_) parameters of our Potts model (Fig. 1a, *Top*). We generated models using both mean-field [2] and Boltzmann machine learning [20] implementations of DCA (mfDCA and bmDCA) to analyze which model was better at predicting functionality. Starting with the *E. coli* WT TEM-1 protein sequence (UniProt ID P62593), variants are computationally evolved through the SEEC model. Specifically, sites to be mutated are chosen at random over the entire protein. For a particular chosen site, a substitution is made based on a conditional probability distribution for mutations that is calculated based on the family model parameters [1] (see *Methods*). Once the simulation of N number of steps is completed, a Hamiltonian, or statistical energy, is calculated for each sequence along the evolutionary trajectory. Using these scores and various other selection parameters discussed further on, key variants are selected for the next phase of *in vivo* experimental testing. In the experimental phase (Fig. 1a, *Bottom*), we obtained synthesized versions of these variants, cloned into an expression vector under the control of an inducible promoter (see *Methods*). *E. coli* clones were grown under the challenge of ampicillin and the optical density was monitored over time. The results from the variant growth assays were used to feedback into the SEEC model for the optimization of computational evolution.

**Fig. 1:**
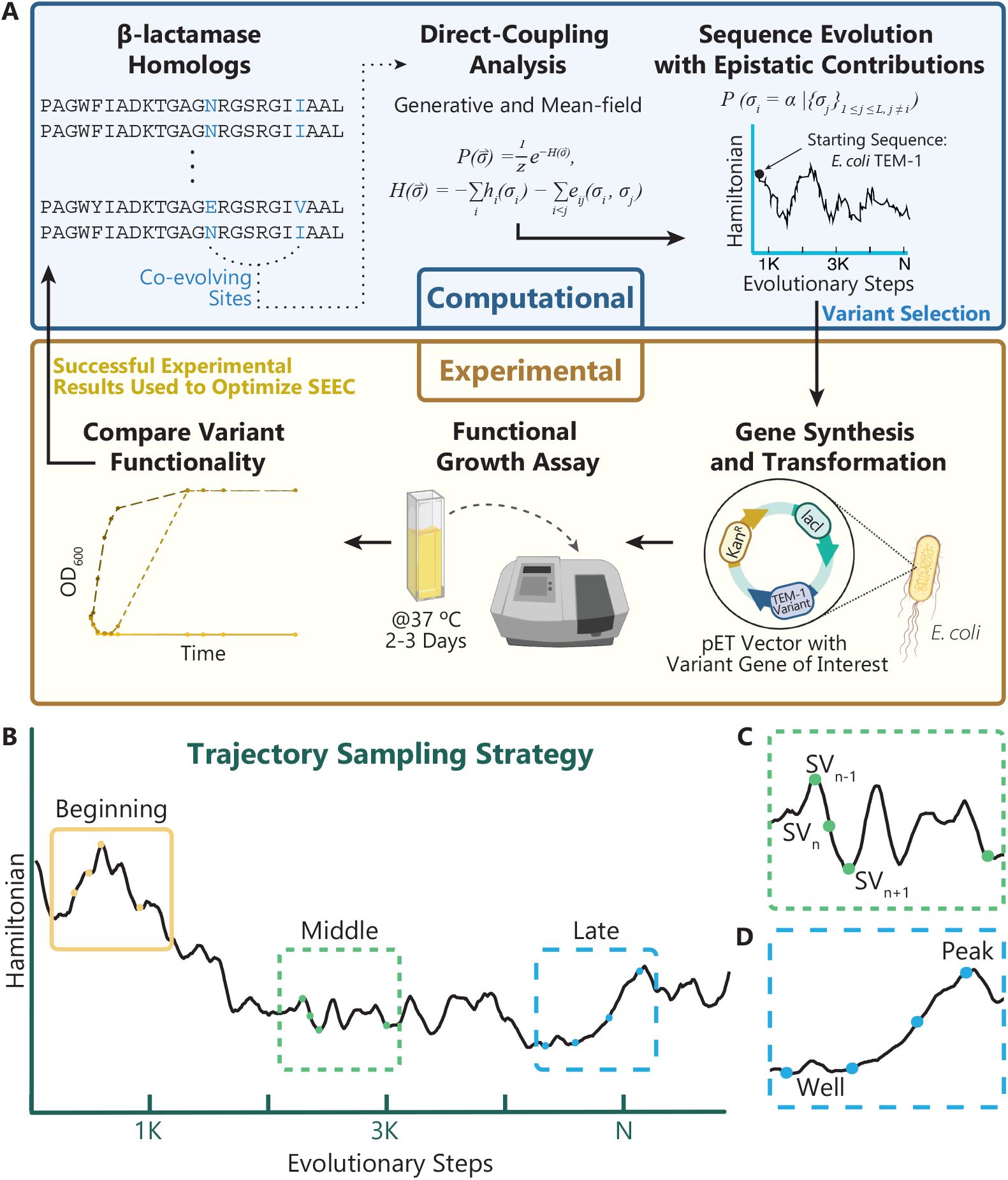
Schematic representation of the computational evolution and experimental validation cycle. (A) The MSA informs the statistical parameters for both the mean-field and generative (Boltzmann machine) DCA models. Starting with the *E. coli* wild type TEM-1, variants are computationally evolved through a process of iterative site mutation over N evolutionary steps (SEEC) and key variants are selected for testing. Each TEM-1 variant is cloned into a pET vector with essential expression and selection genes. Once transformed, the cultures are tested in the presence of antibiotics. The results of variant functionality are used as feedback for the optimization of the computational evolution. (B) Strategy for sampling variants across evolutionary trajectories. To completely sample the evolution, three areas are established: the beginning, middle, and late portions of the trajectory. Within a given area, additional selection attributes include (C) sampling sequential evolutionary steps and (D) selecting variants with increasingly negative Hamiltonian scores, described as *wells* of sequence space, and variants with more positive scores, sampled from *peaks* along the evolutionary trajectory.

We employed three strategies for choosing variants from our evolutionary trajectories. First, in order to sample the evolutionary trajectory, we established three areas for variant selection: the beginning, middle, and late portions of the trajectory (Fig. 1b). In doing this, we set to explore if there existed a threshold for the number of changes one single protein could undergo while still retaining the original function. Secondly, we targeted sets of variants that fell sequentially across evolutionary steps (Fig. 1c). These sets would include triplets of protein sequences that differed only by a single point mutation between each step. Our hypothesis is that SEEC recapitulates neutral evolution, in that fitness might be compromised in some steps, but as long as the gene remains viable, later changes can improve fitness *in vivo*; this strategy for variant exploration aims to assess the presence of this feature. Lastly, we analyzed variants across a range of Hamiltonian scores: many with favorable scores that were increasingly negative, often found in wells within the Hamiltonian trajectory, some with more positive scores, found in peaks along the trajectory, and other sequences with average scores picked from areas between wells and peaks (Fig. 1d). There is evidence that with optimized simulations and generative models that utilize a Markov search process, you can produce biologically active sequences [21, 22]. It follows then that with such models local changes in Hamiltonian energy that result from the simulation parameters could also be predictive of functionality [23]. In using these strategies, we aimed to further understand the multifaceted predictive power of simulated evolution via SEEC.

### SEEC-amino acid struggles to produce functional variants

Phase I of variant testing began with the original SEEC algorithm, where any position across the Pfam domain model (N = 202) could be mutated to either an amino acid or gap. Here, we refer to this version of the model as “SEEC-amino acid” (SEEC-aa). The first computational evolution was performed using parameters inferred using bmDCA (Supplementary Fig. 1) due to its generative potential and correspondence to our previous studies [1] wherein our findings utilized the Boltzmann machine learning implementation of DCA (Fig. 2a). The simulated proteins trended very quickly away from the WT Hamiltonian score (*H* ≈ -550), so we chose our three key areas within the first half of the overall trajectory and utilized the selection strategies specified in Fig. 1b-d. To explore the potential benefits of alternative statistical models, we performed an additional simulation using parameters inferred via mean-field DCA (Fig. 2b, Supplementary Fig. 1). In our previous work, we found that this model resulted in output sequences with lower Hamiltonian energies, which could be indicative of improved fitness [1, 9, 15, 18, 19]. Correspondingly, we found that the mfDCA trajectory quickly trended downwards towards lower Hamiltonian energies compared to the bmDCA simulation. For comparison, key variants were selected that shared similar percent identity as those selected from the bmDCA trajectory. All SEEC-aa variant sequences can be found in (Supplementary Table 1).

**Fig. 2:**
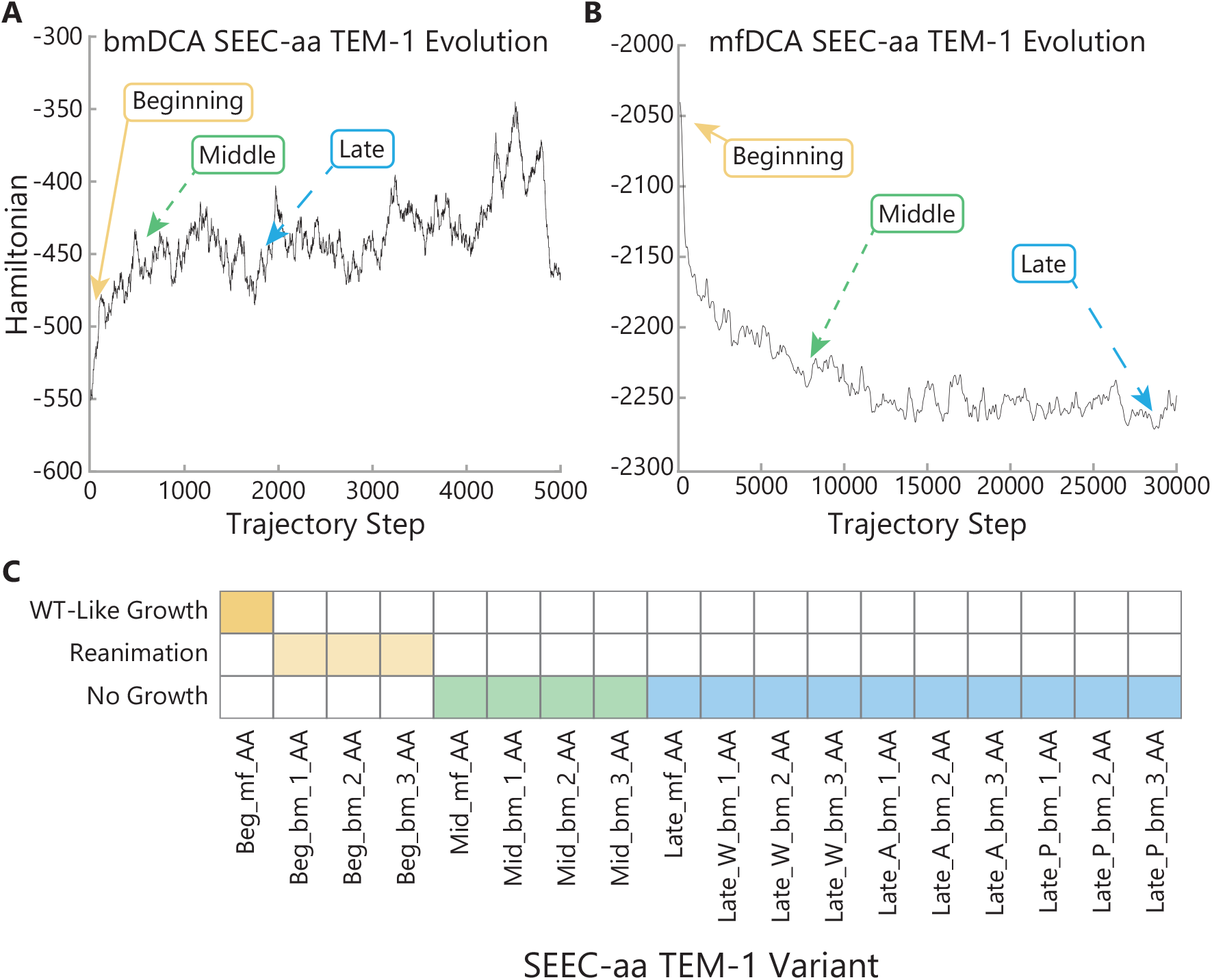
Amino acid Sequence Evolution with Epistatic Contributions (SEEC-aa) Phase I variant selection and functional results. (A) Boltzmann machine learning (*bm*) evolutionary trajectory and areas of variant selection. (B) Mean-field (*mf*) evolutionary trajectory and areas of variant selection. (C) Results of variant functional growth assays. At MIC of ampicillin, the beginning *mf* variant grew comparable to WT TEM-1, the beginning *bm* variants exhibited a delayed function termed *reanimation*, and all other variants demonstrated no resistance to the antibiotics. W = Well, A = Average, and P = Peak.

When cultures containing cells transformed with these variants were grown in the presence of ampicillin, we found that a majority of sequences could not survive even at the minimal inhibitory concentration (MIC) of 5 *μ*g/mL (Fig. 2c, Supplementary Fig. 2). From this, we concluded that for the bmDCA evolution, the Hamiltonian sequence energies that trended in a positive direction quickly left functional space. Even though all tested variants had Hamiltonian scores that were still in the range of the native family sequences (Supplementary Fig. 3), they came from a trajectory that suffered from an initial burst in Hamiltonian score, likely signifying that all subsequent variants would be outside of the original WT functional area. Bacterial cultures expressing the bmDCA beginning variants with 2, 3, and 4 amino acid changes experienced immediate death and only exhibited function and growth after a delayed period; we have termed this growth phenotype as *reanimation* (Fig. 2c). However, the mfDCA beginning variant with 3 amino acid mutations did not exhibit this same phenotype – instead, these cultures survived on par with the WT at both MIC and standard levels of ampicillin (50 *μ*g/mL) (Supplementary Fig. 4). We attribute this success to the fact that the evolutionary trajectory trended downwards into a more favorable, negative Hamiltonian space. Still, not all the mfDCA variants were functional, and this short-coming could be due to the fact that despite trending favorably, the middle and late variants’ scores fell outside the family Hamiltonian distribution (Supplementary Fig. 3). Overall, these results suggested that optimizing the simulation parameters to modulate the initial direction of the trajectory could maximize the number of functional variants, a strategy recently shown to be important by other groups [21, 24].

### Inclusion of simulation restrictions optimizes SEEC functional phenotypes

Using the information we learned from the first round of testing, we adjusted relevant aspects of the computational process for further optimization. One concern we identified was that the evolutionary model parameters were inferred from the Pfam domain family, which only contained 202 out of 263 total amino acids (Supplementary Fig. 1c, see *Methods*). Since mutations acquired during the simulation only occurred in this area, there was a potential that the evolved sequence might have lost compatibility with the retained WT portions outside of the domain. To address this issue, we chose to create a better contextualized model by using an MSA queried on the entire *E. coli* TEM-1 sequence without the signal peptide (Supplementary Fig. 5c).

In doing so, we increased the specificity of our MSA for TEM-1 like proteins, but also decreased overall diversity as the effective number of sequences changed from 3,834 to 1,152. The impact of this greater specificity can be seen in the comparison between the Pfam contact maps (Supplementary Fig. 1) where the number of true positives is greater than the number of hits for the TEM-1 whole protein contact maps (Supplementary Fig. 5). Although our current hypothesis is that our model can produce functional proteins, they might not be optimized for a specific cellular context. Increasing sequence diversity is known to improve structure prediction, but granting the simulation access to broader sequence space could be pushing simulated proteins towards different realms of taxonomic evolution that are discordant with the organismal context of the input sequence [15, 25]. While all the *β*-lactamase family members are catalyzing essentially the same chemical reactions, the proteins themselves are acting in different biological contexts. The physiology of this process includes protein-protein interactions, optimal growth temperatures, expression/translation regulation, and so on - all which require specific context provided by the host organism. Therefore, considering the taxonomic context of an evolved gene influences the potential of not just its biochemical functionality but its physiological implications as well [26, 27]. We reasoned that funneling the explored sequence space to that which corresponded to the specific physiological context of *E. coli* would enhance the goal of pursing evolutionary trajectories with functional, novel proteins. In addition to changes made to the input aligned sequences, we also made adjustments to the SEEC algorithm to address obstacles experienced in the first iteration of experimental trials. For instance, we modified the mutation criterion so that the algorithm could no longer introduce gaps or select gaps for mutation during the simulation which would prevent shortened or lengthened output proteins. Additionally, we again selected amino acid substitutions based on a conditional probability distribution, but this time, only residues that were accessible via a single nucleotide change could be accepted. Bisardi et al. found that in making these changes, simulations from a similar model better correlated with *in vivo* functional data [24]. These changes helped further establish the biological relevance of the SEEC evolutionary model (hereafter referred to as SEEC-nucleotide or SEEC-nt) with the goal of improving functional variant output.

The final improvement came from optimizing the simulations themselves by modulating the model *selection temperature* [12, 21]. During the initialization of the evolutionary simulations, there exists the opportunity to regulate the family couplings (*e*_*ij*_) and local fields (*h*_*i*_) statistics with selection temperature (*T*) from Eq. 3 (see *Methods*). Using the Hamiltonian from Eq. 2 as a representation of statistical energy, adjusting the temperature of these parameters learned from the protein family serves as a method to adjust the energetic exploration of the mutational space. During phase I, we did not select specific temperatures for each simulation as both mfDCA and bmDCA inferred models were ran at temperature = 1. From these experiments, we saw improved results with the lower sequence energy trends from the mfDCA SEEC-aa variants. Moving into Phase II, we applied the same concept to our bmDCA model to sample at temperatures less than one for resulting sequence trajectories trending towards favorable Hamiltonian values.

### SEEC-nucleotide produces variants with improved functionality

Using SEEC-nt, we again ran simulations using parameters inferred from both mfDCA and bmDCA and generated 5000-step evolutionary trajectories starting from the WT *E. coli* TEM-1 sequence. In contrast to error-prone PCR mutagenesis or saturation mutagenesis, our global model informs the propensity for a change at the current step by all of the mutations that have come before in the evolutionary simulation. From these simulations, following the same strategies described earlier, we chose thirteen variants using both mfDCA and bmDCA to infer parameters (Supplementary Table 2). For these trajectories, the late variants that had evolved the most had acquired 34 point mutations for Late mf 3 NT and 58 point mutations for Late bm 3 NT compared to the WT TEM-1. When these late variants were queried on BLASTp [28], the top hits were to the *Pseudomonas aeruginosa* TEM-136 that has 33 different point mutations from Late mf 3 NT and a *Citrobacter freundii* class A *β*-lactamase that has 48 different point mutations from Late bm 3 NT.

This further supports that these variants are exploring novel sequence space rather than becoming another extant protein. Figure 3a details the accumulation of mutations for the variants chosen from the bmDCA evolutionary trajectory. While there are some mutation steps that are reversions back to the WT sequence (blue dots), most mutations carry the protein into novel sequence space with the late variants having close to 60 different substitutions. Note that variants Beg bm 4 through 6 were sampled from the earliest part of the trajectory, and as such have the least accumulated mutations. We selected these variants in order to make direct comparisons to the mfDCA sequences with equal numbers of mutations. Panels b and d on Fig. 3 contain the trajectories with points and areas marked for the variants chosen for testing. For the mfDCA model, due to the changes made in phase II, we had to increase the simulation temperature to 1.5 to see noticeable changes in the protein evolution (Fig. 3b, Supplementary Fig. 6, see *Methods*). Here, we can see that the trajectory favorably becomes more negative, and the variants’ Hamiltonian scores relative to the simulation temperature remain similar to WT and remain within the distribution of scores for the family (Supplementary Fig. 7). Initially we tested *in vivo* functionality of these variants at the MIC for ampicillin (5 *μ*g/mL) and found that growth was not challenged at all, so we raised the ampicillin concentration to 50 *μ*g/mL (Fig. 3c). Final samples were collected and Sanger sequenced; quality chromatograms covering the entire gene were obtained for the majority of the samples, enabling us to conclude that our sequences of interest remained intact during the experiment. Also, no significant population with additional compensatory mutations acquired during the assay can account for the observed phenotype (see Supplemental Information for Chromatograms). For the mfDCA SEEC-nt variants, even this level of 50 *μ*g/mL ampicillin was not a challenge as they all grew at rates on par and sometimes better than WT (Supplementary Fig. 8).

**Fig. 3:**
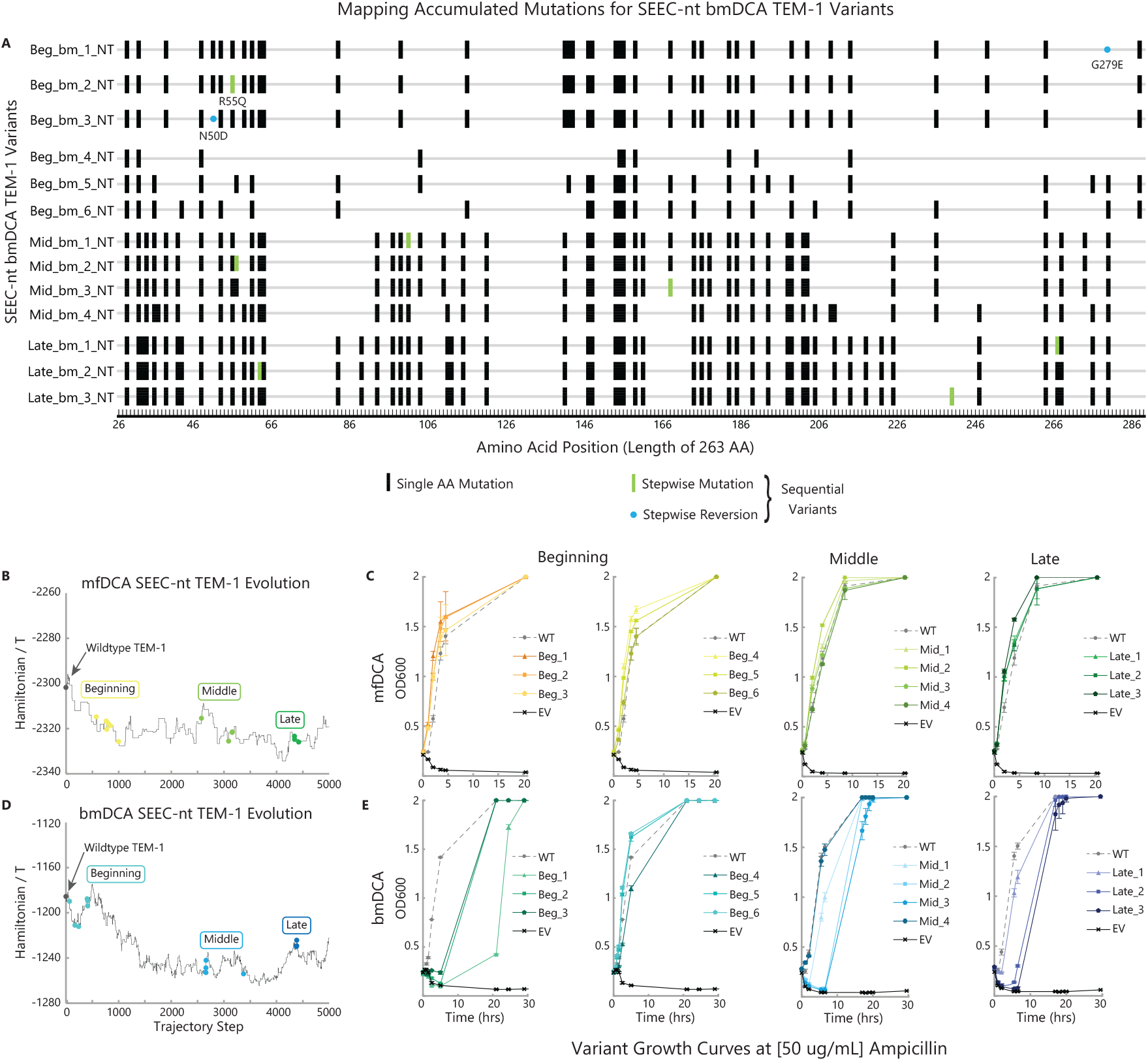
Nucleotide Sequence Evolution with Epistatic Contributions (SEEC-nt) Phase II variant selection and function results. (A) Sequence diagram of variant mutations for the SEEC-nt bmDCA TEM-1 variants. Singular change between sequential variants displayed as green bars for mutations or blue circles for reversions. (B) Mean-field (*mf*) at T = 1.5 and (D) Boltzmann machine learning (*bm*) at T = 0.75 evolutionary trajectories and areas of variant selection. (C) *E. coli* growth curves for variants picked from the mfDCA and (E) bmDCA model SEEC-nt trajectories. Cultures were grown in 50 *μ*g/mL ampicillin. Data points are the mean of 3 experimental replicates and error bars represent standard deviations. EV = Empty Vector.

Likewise, the bmDCA trajectory also becomes favorably negative, however most of the middle and late variants’ scores are outside of the family range (Fig. 3d, Supplementary Fig. 7). As expected, the simulation temperature for the bmDCA model had to be modulated to achieve a favorable Hamiltonian trend, so this simulation was run at T = 0.75 (Supplementary Fig. 9). Interestingly, despite requiring a low selection temperature, the bmDCA simulations still explored vaster sequence space than the mfDCA simulation ran at twice the temperature (compare Percent ID between Supplementary Fig. 9 and Supplementary Fig. 6). When tested at 50 *μ*g/mL ampicillin, select beginning and middle variants (Beg bm 5 NT, Beg bm 6 NT, and Mid bm 4 NT) grew on par or faster than WT, while all other variants facilitated only slow growth or reanimation (Fig. 3e, Supplementary Fig. 10). When considering the difference in results between the SEEC-aa and the SEEC-nt variants, the beginning variants for bmDCA SEEC-aa only had 2, 3, and 4 mutations that nevertheless resulted in an impairment of function and delayed growth potential leading to reanimation. Correspondingly, while there exists the threshold model that random mutations would impact stability and not function directly, once many of said mutations accumulate, their effect will exponentially decrease the protein’s overall fitness [29]. Specifically within TEM-1, Bershtein et al. found that the synergistic accumulation of 8 or more random mutations more than exponentially diminished fitness [30]. Thus the fact that bmDCA SEEC-nt generated a variant that retained WT-like activity, such as Mid bm 4 NT with 47 substitutions, is a remarkable result. The potential number of sequences restricted by mutation in nucleotide space is approximately 9^47^ positional changes, hence this variant exists in a possibility space that exceeds the total number of water molecules on the earth. Therefore, finding this sequence by chance is not plausible.

### SEEC-nt informed by mfDCA evolves variants with enhanced enzymatic activity

To glean further insights on the difference in functionality among phase II variants, we further analyzed the sequence trends and their behavior at even more challenging levels of antibiotics. In Fig. 4, we can see the expanded SEEC-nt trajectories with individual variants pinpointed along the simulation (Fig. 4a). To push the functional boundaries of these variants, we tested them all in cultures containing a challenging level of 100 *μ*g/mL ampicillin (Fig. 4b). In quantifying the different rates of growth for the variants and WT protein, we calculated the ratio between the mean optical density of the experimental samples and their positive controls and proceeded to compare the rationalized growth between each variant and the WT (detailed in equations 5 and 6, see *Methods* for details). By performing this analysis, we found that many variants exhibited multiple folds change in activity with positive peaks illustrating enhanced growth during exponential phase while fold growth less than one represents sub-optimal function when compared back to WT. Surprisingly, every mfDCA variant outperformed the antibiotic resistance activity of the WT protein by several fold (Fig. 4b, *mfDCA Fold Growth* and Supplementary Fig. 11). On the whole, for this mfDCA simulation, almost every variant achieved an increase in fold growth when compared to the activity of variants from the previous section of the trajectory. Remarkably, the best fold growth was seen from Late mf 2 NT (Fig. 4b) with over a 10-fold increase in the rate of culture growth during the exponential phase of the growth assay. Therefore, even with an increasing number of accumulated mutations, the global downward trend of the Hamiltonian*/T* mfDCA SEEC-nt trajectory is highly predictive of improved variant functionality.

**Fig. 4:**
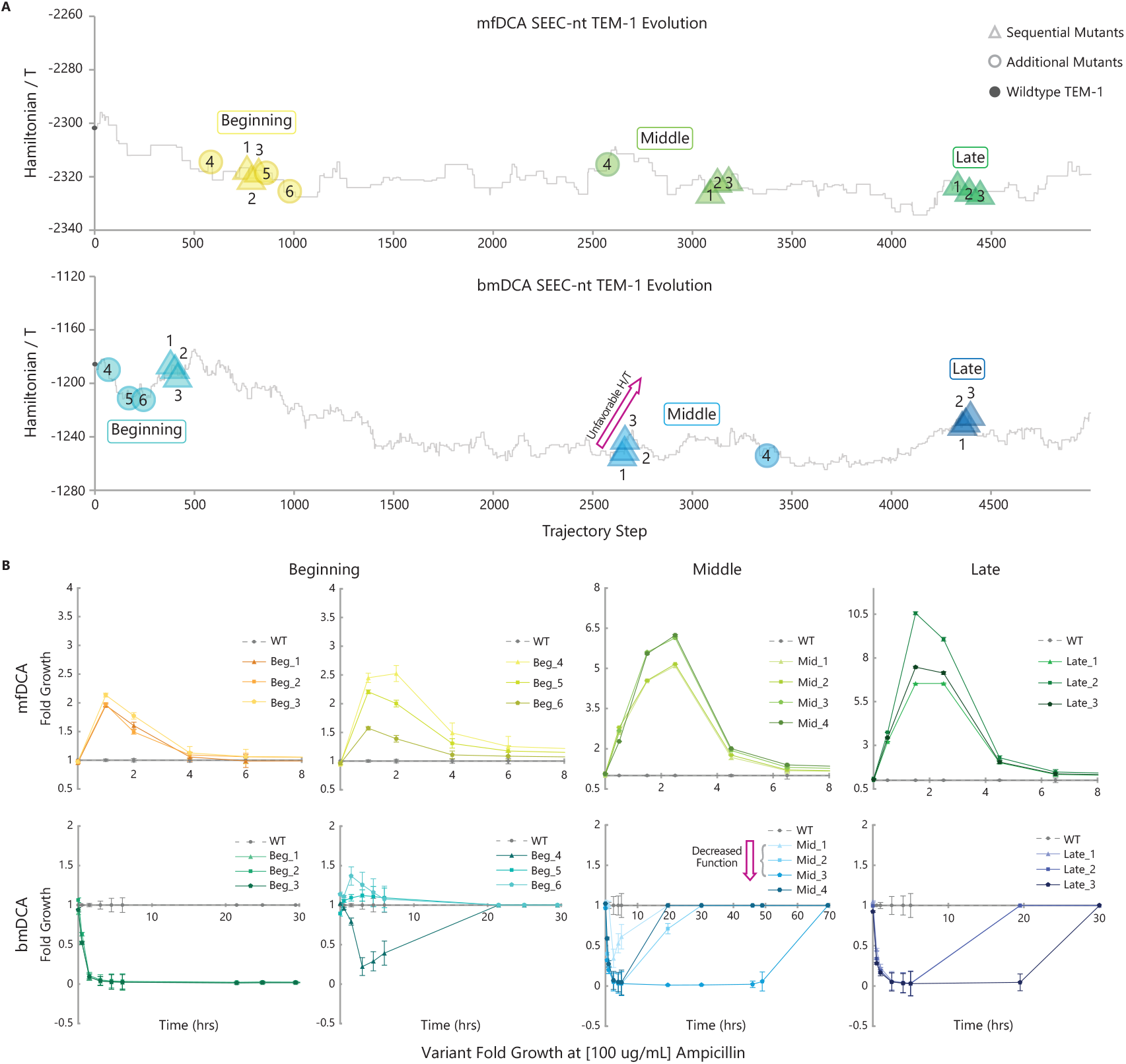
SEEC-nt Phase II variant fold change in function compared to WT *E. coli* TEM-1. (A) Mean-field (T = 1.5) and Boltzmann machine learning (T = 0.75) expanded evolutionary trajectories and areas of variant selection. Points of variant selection from the trajectories are indicated with corresponding colors and shapes based on area and mutant type. The purple arrow highlights the pattern of the Hamiltonian divided by temperature score becoming increasingly unfavorable for the middle **bmDCA** sequential variants. (B) Fold growth graphs for variants picked from the mfDCA and bmDCA model SEEC-nt trajectories. Cultures were grown in 100 *μ*g/mL ampicillin. Within the fold graph for the middle **bmDCA** variants, the purple arrow emphasizes the pattern from panel a where the Hamiltonian score becomes unfavorable, a decrease in variant functionality follows. Data points are the mean of 3 experimental replicates, then normalized to the mean of 3 positive controls detailed in Eq. 5, then normalized to the WT rationalized growth from Eq. 6 (see *Methods*). The error bars represent the addition of propagated errors from the standard deviations of the measured samples.

### SEEC-nt informed by bmDCA highlights predictive power of the local Hamiltonian context

Beyond the global trend of the trajectory becoming more negative, we find that crucial, predictive patterns also occur in the *local* context. For example, in the bmDCA simulation, the sequential middle variants increase in Hamiltonian*/T* scores in a step-wise fashion, gradually becoming less favorable (Fig. 4a, *Bottom*). In the case of these variants, as the Hamiltonian becomes increasingly unfavorable, we see a decrease in ampicillin resistance as it takes each sequential mutant a longer period of time to reanimate (magenta arrows, Fig. 4). A similar pattern can be said of the bmDCA late variants as well. The same phenomenon occurs, but in the opposite direction, for the bmDCA beginning sequential variants. These sequences, albeit selected from an early, higher energy portion of the trajectory, are sequentially moving towards a local *well* of Hamiltonian energy. While these variants could not survive at this challenging level of ampicillin (100 *μ*g/mL) (Fig. 4b, *bmDCA Fold Growth* and Supplementary Fig. 12), in Fig. 3 where the concentration was only 50 *μ*g/mL, we again see a step-wise pattern in function. This time, however, the final sequential variant with the lowest Hamiltonian, Beg bm 3 NT, retains the best functionality between the three proteins and can grow to saturation the quickest, followed by the second and then the first beginning variant.

While the global trend of the trajectory is meaningful, it is not enough to predict individual variant functionality. If the global movement were sufficient, then each variant from a negative trending trajectory would be functional. However, our data shows this is not the case, as demonstrated with the difference in function between the bmDCA middle and late variants when compared to Beg bm 5 NT and Beg bm 6 NT. Despite having more positive Hamiltonians, these two variants outperform their later variant counterparts, even in the most challenging levels of ampicillin. This effect is not easily explained by the early positioning of these variants in the trajectory, as each has already accumulated 26 mutations. At the same time, in Fig. 4 we have two examples in which sequential variants that head towards a local peak coincide with decreased functionality (middle and late bmDCA variants), and one example in Fig. 3 in which sequential variants that head towards a local well coincide with increased functionality (beginning bmDCA variants), indicating that it is in fact these local Hamiltonian trends that are a more reliable marker for optimizing functionality.

### Boltzmann machine learning model allows SEEC-nt to explore greater sequence space

Our results support the idea that there is a clear difference between the functionality of our variants chosen from models inferred using mean-field versus Boltzmann machine learning implementations of DCA. While both models produced functional variants, mfDCA consistently produced variants that outperformed the WT controls. Part of the reason for this could be that the variants chosen from the bmDCA model explore more sequence space, even with the constraints provided by making mutations in nucleotide space. The logos in Fig. 5a, b depict the frequencies of amino acids found at each site in the two SEEC-nt variant pools that we experimentally synthesized and assayed. At several positions, highlighted by orange dots, the bmDCA model samples more amino acids (Fig. 5). This is especially meaningful because the mfDCA group has more variants with a substitution at these sites (Supplementary Fig. 13a, b). We also looked at sites that are mutated at least once in both trajectories, compared the substitutions made for the mfDCA and bmDCA trajectories (Supplementary Fig. 14) and noted the amino acids that were common to both or unique to either of the models. As with the logos, this analysis similarly reveals that sequence space exploration is more extensive in the bmDCA model, as the number of unique amino acids are overwhelmingly greater in the bmDCA-based trajectory (compare orange versus green). Our chosen variants are representative of the simulation as a whole, as we see the same phenomenon occurring there as well (Supplementary Fig. 14, *Bottom*).

**Fig. 5:**
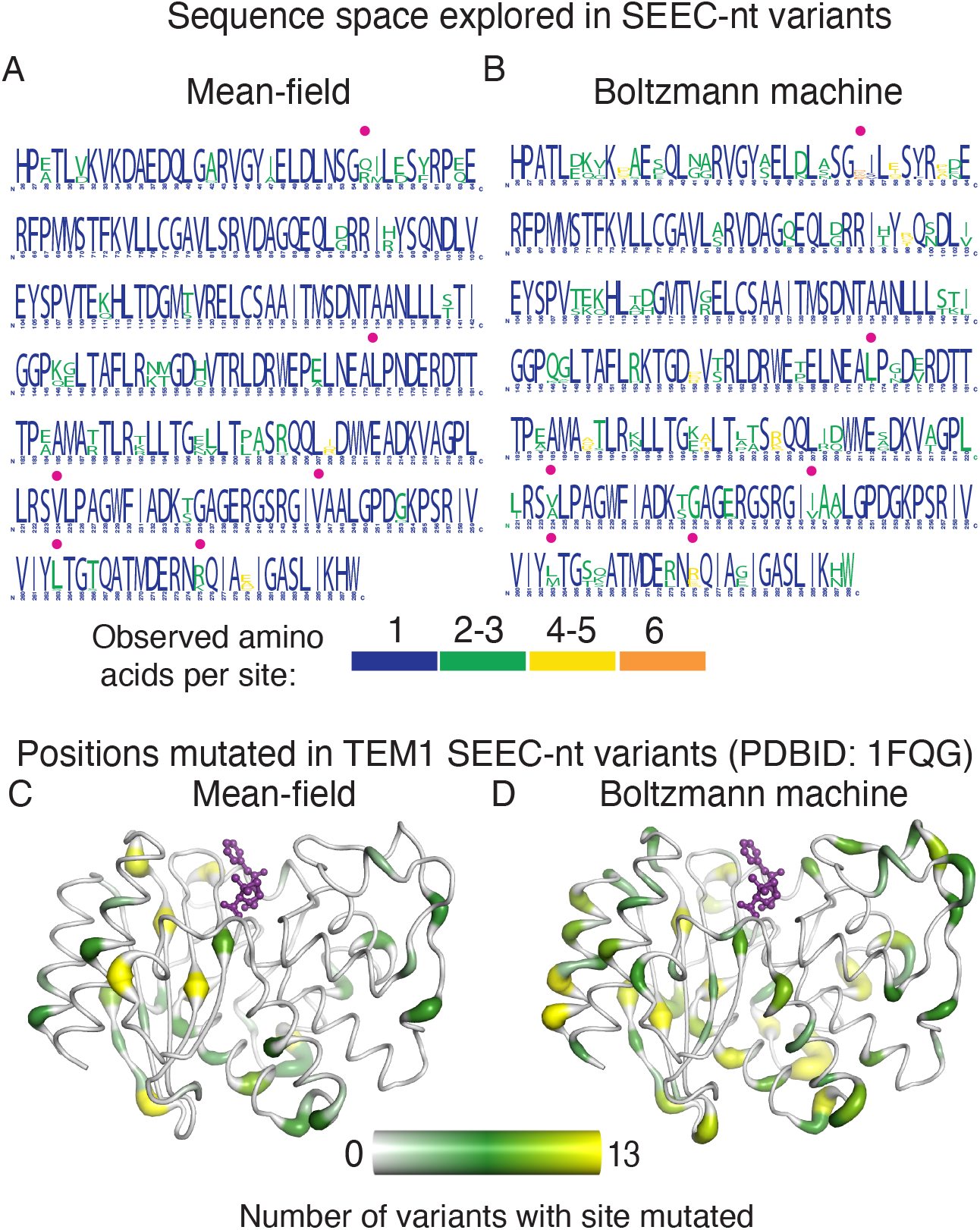
Comparison of sequence space exploration within mean-field (*mf*) and Boltzmann machine learning (*bm*) Phase II variants. Logo of amino acid frequencies across the 26 variants tested, 13 from *mf* (A) and 13 from *bm* (B) models. Orange dots highlight examples of positions which showed more amino acid diversity in the *bm* group of variants, despite being mutated more often in the *mf* group of variants. Logo generated as a frequency plot with small sample correction using https://weblogo.berkeley.edu/logo.cgi. Colors indicate groups of similar amino acids. (C) and (D) The frequencies of each position being mutated across the variants derived from *mf* (C) and *bm* (D) models are mapped onto the TEM-1 structure (PDBID: 1FQG) using a color gradient and putty thickness. Penicillin D, covalently linked to the nucleophilic Serine residue, is depicted in purple ball and sticks and highlights the active site region.

To further demonstrate the greater freedom of bmDCA sequence space exploration, we analyzed the amino acid frequencies for each site across the 5K variants generated by each simulation (Supplementary Fig. 15a). In the heat maps, columns with a dark red rectangle (frequency = 1) in a dark background (frequency = 0) indicate conserved sites that never changed throughout the simulation. Comparison of the two heat maps reveals that the bmDCA model produces far more sites in between these two extremes, signifying more sequence space exploration. Importantly, a scatter plot of the amino acid frequencies from both trajectories highlights that often, a position is conserved during the mfDCA trajectory (*X* = 1) but mutated during the bmDCA (*Y <* 1, 158 points, purple oval); it is rare, however, to see this in the other direction (4 points, light blue oval, Supplementary Fig. 15b). Do all amino acids get treated the same by both models, or are some amino acids more substituted in one model versus the other? To address this question, we compared amino acid frequencies for mfDCA versus bmDCA models by calculating the correlation coefficients for each set of amino acid frequencies across all sites (Supplementary Fig. 15c). In panel d, this calculation is shown for the 5K trajectory sequences as well as the subset that were chosen for experimental testing. For the trajectory data (x-axis), all of the Pearson correlations of these amino acid frequency vectors for the mfDCA and bmDCA models are over 0.85 for all amino acids except Asn, which is ∼ 0.70. The correlations are even higher for the chosen variants (y-axis). The amino acids that are most similar between the trajectories for the two inferred models (that is, close to the diagonal) are Cysteine, Tyrosine and Tryptophan, while the residues that change the most are Asparagine and Glutamine. Overall, the scatter plot of these correlations reveals that i) positional amino acid frequencies for each residue type are are different across the mfDCA and bmDCA models, ii) the correspondence is high (*R* = 0.88) between the amino acid frequencies of the whole trajectory and the mutants chosen for experimental testing, and iii) points fall on either side of the line of unity indicating minimal systematic bias, confirming that our experimentally tested cohort are representative of the trajectory.

### Substitutions for both mfDCA and bmDCA models are spread across the structure while avoiding sites with vital functional roles

In addition to looking at the differences in sequence space exploration between the two models, we also asked whether the locations within the structure of the substituted positions differed significantly. Localization of mutation sites on the three-dimensional structure of TEM-1 *β*-lactamase reveals that mutations from both models are distributed across the entire structure except for two regions: one buried helix (E64-L81), which houses essential catalytic residue Ser70 and is nestled between a second conserved region, a helix turn helix (E121-L139)(orange regions in Fig. 5c, d and Supplementary Fig. 13a, b). Despite the conclusion from various computational mutation models predicting solvent exposed sites to be more robust to mutation than buried sites [31, 32, 33], in our variant pool there is no connection between the propensity for a site to be mutated and the degree of solvent exposure of that position in the structure (Supplementary Fig. 13c, d). There is also conserved information about the active sites and correspondingly, the variants retain most of the active site areas. Specifically, the active site residues, Ser70, Gln166 and Asn170 are conserved in both variant cohorts (Supplementary Fig. 13a, b). Even though these critical residues are never mutated, some mutated residues are within 8Å of these active site residues (red dashes, Supplementary Fig. 16a, b). Interestingly, the majority of the substitutions change to similar amino acids. In the cases where this is not true, the bmDCA pool has more variants with non-conservative substitutions (see bold rows, Table 1). Thus, compensatory changes can occur even within contact distance from sensitive active site residues.

**Table 1:** Mutations to residues nearby active site

| position | WT residue | mutated residue | MF | BM |
| --- | --- | --- | --- | --- |
| **168** | **Glu** | **Ala** | **2** | - |
| 173 | Ile | Leu | 13 | - |
| 235 | Ser | Thr | 8 | 8 |
| 103 | Val | Ile | - | 8 |
| **167** | **Pro** | **Thr and Ala** | - | **8** |
| 239 | Glu | Asp | - | 1 |

Clearly there is general conservation in the active site region, and while this is most likely constrained by catalytic requirements, we also investigated the role of other biophysical constraints such as protein-protein interactions. *β*-lactamase binding protein (BLIP) and BLIPII are proteins that abrogate the catalytic activity of *β*-lactamases by binding the conserved helix-loop-helix region (red region, Supplementary Fig. 16c, d) of class A *β*-lactamases and inserting loops into the active site [34, 35]. Biochemical studies have identified the key residues mediating the inhibitory binding activity of these two proteins [36]. Mutations within our experimentally tested cohort are almost nonexistent among the BLIP and BLIPII binding hot spot positions in TEM-1: E104,L102, Y105, P107, K111, and M129 (Supplementary Fig. 16c, d). Taken together, these observations highlight the fact that SEEC-nt does not merely target positions that are “low hanging fruit”, or changes that would be obviously non-disruptive, like surface residues. At the same time, the preserved regions are likely to be essential across the family for either structure, catalytic efficiency or even protein-protein interactions; this all demonstrates the benefit of a model informed by family statistics.

## Discussion

In this work we provide evidence on how SEEC, an epistatic evolutionary model, leads to sequence changes that preserve function. We observe this result not only for a few mutations, but many across the entire protein. As we saw with the initial SEEC-aa trials, the potential for a handful of mutations disrupting function is evident, hence the number of mutations acquired with SEEC-nt that continue to retain functionality is significant. In the case of the mean-field DCA model, we created variants with sequential changes that still preserved function, and in every case, resulted in improved antibiotic resistance capacity when compared to WT. These results further demonstrate that the SEEC model recapitulates neutral evolution, and that with mfDCA-inferred parameters, each sampled evolutionary step retained fitness and viability *in vivo*. This model presents a relatively faithful reflection of neutral evolution. Through these simulations, we observe how the proteins traverse sequence space, how their functions are impacted by various mutations, and the overarching, cyclical nature of evolution. Over time, sequences fluctuate for better or worse functional activity. These resulting proteins represent the culmination of thousands of evolutionary events, and in the end, the number of functional mutations after this magnitude of potential changes is remarkable. Although we have noticed in previous work that SEEC had convincing statistical features for evolutionary models, we now provide evidence of plausible evolutionary trajectories that lead to functional phenotypes *in vivo*.

When comparing the two versions of the statistical model, we aimed to fairly evaluate simulated sequences by selecting a handful of variants from both models that had accumulated the same number of mutations during their respective simulations. Comparatively, SEEC-nt informed by bmDCA headed into novel sequence space faster than the mfDCA simulation, so variants had to be selected from the beginning portion of the trajectory to match the percent identity of the mfDCA selected variants (Supplementary Table 3). In every comparison between variants with equivalent numbers of changes, mfDCA always led to more fit, functional variants than their bmDCA counterpart, even at the most challenging level of ampicillin. Concurrently, we found that mfDCA informed SEEC-nt consistently resulted in functional sequences that all had better antibiotic resistance activity than WT. We envision that these mfDCA-inferred parameters could represent an evolutionary scenario in which the selection pressure is higher, thereby restricting the evolution to variants with increased fitness. On the other hand, the bmDCA simulations also resulted in many functional proteins, just some with better and some with worse antibiotic resistance than WT. Here, the generative model obtained with bmDCA could represent an environment with a lower selective pressure, allowing for more freedom to explore sequence space. These observations highlight the fact that SEEC, as a platform to model and learn about evolution can incorporate epistatic constraints from any source including experimental data. In addition, the epistatic relationships need not be limited to second order: higher order interactions have been found through both computational [37, 38, 39] and experimental methods [40, 41, 42].

To further decipher the difference in functional activity between the two versions of the model, we looked into elements that could inform the propensity for mutation across different regions, such as catalytic regions, specific protein-protein interfaces, and how they impact which substitutions will be tolerated at specific sites. We found that the BLIP and BLIPII binding hotspots were conserved among our tested SEEC-nt variants for both mfDCA and bmDCA models. As an enzymatic inhibitor, it would make evolutionary sense that BLIPs would target a region necessary for catalytic function. We speculate that inserting multiple mutations in this region, as is the case in our bmDCA mutants, has diminished catalytic function. Simultaneous mutations at multiple sites in this region have been shown to slightly harm catalytic efficiency [43]. Given that there are more of these positions mutated in the bmDCA pool, this might explain the reduced activity of these variants in comparison to their mfDCA counterparts. Further studies exploring the BLIPs’ binding activity of SEEC-nt variants could clarify the picture of evolutionary and physiological constraints on the explored mutational space in our simulations.

The future of the SEEC model includes exploring the prospects of neofunctionalization; by steering evolution into specific new fitness minima of the sequence-function landscape, novel functions and contexts could be investigated. While building functional models is important work, we can also apply this knowledge of evolutionary constraints now to other subjects. There exists the potential to trace functional pathways for the evolution of viruses using the existing sequence space. With this data, we could study viruses such as HIV or SARS-CoV-2, and potentially aid the development of vaccines or therapeutics for future variants before their arrival. Awareness of this virus fitness landscape allows us to utilize proactive design against the most probable variants [44, 45, 46, 47]. This knowledge could grant us the ability to target specific strains before they overtake the population at large in addition to studying the dynamics of the sequence evolutionary change. In addition, one can study how the genome adapts in the presence of SEEC-modified essential genes; the tunability of the function based on local Hamiltonian fluctuations might make new compensatory pathways available that are not accessible when *β*-lactamase activity is completely abrogated. [48]. This is more consistent with the phenomenon of gradual genetic change. The ability to use sequence-based computational models of evolution such as SEEC will continue to provide us better insight into the process of neutral evolution, ancestral reconstruction of sequences, novel protein-design applications, as well as critical predictive power regarding pathogen landscapes during unrelenting epidemics.

## Methods

### Input Sequence Datasets

*E. coli* TEM-1 is a member of the *β*-lactamase2 domain family (entry ID PF13354) in the Pfam database [49]. To test the ability of SEEC-aa to output functional variants of TEM-1, an MSA of homologous UniProtKB database sequences was obtained from the Pfam database. Any homologues with continuous stretches of gaps totaling greater than 5% of the model length (N = 202) were excluded from the final alignment, which came to 15,495 UniProtKB sequences. After reweighting sequences with 80% or greater sequence identity using a pseudocount of 0.5, the effective number of sequences was ∼ 3, 834. For the second phase of model testing, we made several changes in order to improve coverage of the protein domain. First, we modified the domain family definition so it included the entire sequence of TEM-1 (minus the signal peptide), in case there was a problem with accumulating mutations in the domain that eventually became incompatible with the upstream and downstream portions (compare Supplementary Fig. 1C and Supplementary Fig. 5C). Second, with this new model (N = 263) we obtained a new MSA by using the TEM-1 sequence as the seed from HMM Build to construct an HMM profile [50, 51]. With this profile, we then utilized HMM search to obtain matches within the UniProt database, including entries in both Trmbl and Swiss Prot [52]. After filtering to 5% continuous gaps and reweighting as before, the effective number of sequences was ∼ 1, 152.

### Parameter Inference and Hamiltonian

The DCA method [2] was then applied to the MSAs discussed before to estimate the direct coupling between all pairwise residues as well as the residue preferences at each position. As described in [2], DCA utilizes maximum entropy modeling to estimate the joint probability distribution of protein sequences 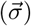:

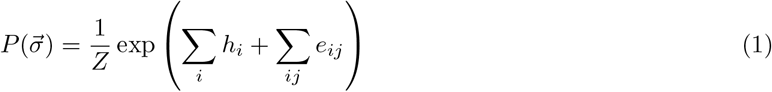

where *Z* is the partition function, the position of residues within the aligned domain or protein sequence are denoted as *i* and *j*, and parameters *e*_*ij*_ and *h*_*i*_ can be numerically inferred. The *e*_*ij*_ parameters quantify the coupling strength for residues *i* and *j* for all possible amino acid occurrence pairs. The amino acid biases for independent positions are captured by the parameter *h*_*i*_. The sums of the *e*_*ij*_ and *h*_*i*_ parameters can be characterized as an energy function, or Hamiltonian (*H*):

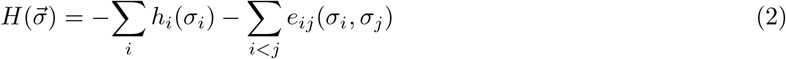

The Hamiltonian represents a statistical energy for a sequence and has been predictive of functional and nonfunctional effects in proteins and RNA [15, 53, 54, 55]. As it so happens, the inference of the exact parameters is an intractable problem, so they are estimated instead using multiple approximations with a variety of complexities and accuracy. In this work we used both the mean-field implementation [2], which places an emphasis on the identification of highly coupled sites and is minimally complex, as well as bmDCA, which unlike mfDCA produces generative protein family models but is computationally expensive [20]. In our previous work introducing the SEEC algorithm, we tested the statistical properties of the evolutionary trajectories using both mfDCA and bmDCA models as input parameters and found that both captured the statistical features found across several theories of neutral evolution [1]. To investigate if both types of models were able to produce functional variants, we performed mean-field and Boltzmann machine learning DCA implementations to infer the model parameters. Original codes for *e*_*ij*_ and *h*_*i*_ parameter inference by DCA were written in MATLAB (The MathWorks, Natick, MA) and previously published at https://github.com/morcoslab/coevolution-compatibility [16] and https://github.com/matteofigliuzzi/bmDCA [20].

### Selection Temperature

One can introduce an additional parameter *T* to restrict the average value of the Hamiltonian being described by Eq. 2. Analogous with treatments in statistical mechanics, this is called a *selection temperature* [12, 21] and it parameterise the DCA distribution as:

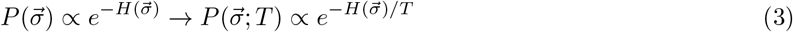

This has the effect of modifying how the overall sampling is restricted. If a change had a probability *p* of happening at a given evolutionary step, now this probability is scaled to *p*^1*/T*^. For larger *T*, this distribution flattens and changes that would not normally be accepted, may now be more probable, increasing the overall Hamiltonian value for the resulting evolutionary trajectory. Conversely, a decrease in selection temperature would restrict changes to only selected mutations with more advantageous Hamiltonian scores, driving the trajectory towards more negative values.

### SEEC-amino acid Algorithm

The *e*_*ij*_ and *h*_*i*_ parameters estimated by DCA, are then used as input for the SEEC-aa evolutionary simulations as previously described [1]. Briefly, this approach chooses a position based on a uniformly distributed random variable and then samples from a conditional probability distribution for all possible amino acids at that site, given the amino acid identities of the rest of the sequence at that step. Importantly, the model of 202 positions excludes 25 residues upstream (not including the signal peptide) and 30 residues downstream of the Pfam domain, and so these positions remain as they are in the WT sequence. For the new sequence 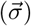, an energy function, or Hamiltonian (*H*), can be calculated from Eq. 2. This new sequence is now the input for the next evolutionary step, and the next position is chosen as before. Finally, a Hamiltonian trajectory can be plotted, which tracks the relative fitness effects of each step in the evolutionary simulation.

### SEEC-nucleotide Algorithm

The SEEC-nt algorithm is similar to the one presented in our previous work [1], but modified to account for mutations at the nucleotide level as well as to prevent insertion and deletion mutations [24]. For this, we track the nucleotide sequence which, with the standard genetic code, translates to the amino acid sequence currently being evolved. At each evolutionary step:

1. One position *i* of the amino acid sequence is chosen by sampling a uniform distribution over all sites that are not gaps.
2. Once chosen, we calculate the probability distribution of the amino acid identity of the site (*α*), conditioned to the rest of the sequence, given by:

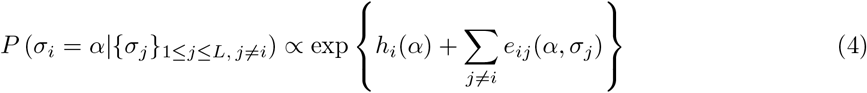

This probability distribution is then sampled. If the resulting sampling selects a gap, it is discarded and the probability distribution at the same site is sampled once again.
3. If, however, an amino acid is sampled, we check if there exists a codon corresponding to this amino acid that is a maximum of 1 nucleotide change from the current codon identity. When this condition is met, the amino acid sequence is updated with the new residue in place, as well as the corresponding codon chosen at random from the possible neighboring codons. When the closest codon differs by 2 or more nucleotides, the site distribution is re-sampled. A site can be sampled up to 100 times before registering as a completed step and leaving both the amino-acid and nucleotide sequence unaltered.

### Variant Selection

To obtain variants, we ran the SEEC-aa and SEEC-nt simulations for several thousand steps under the bmDCA or mfDCA model parameters; we chose a sampling of variants from each trajectory for experimental testing based on the strategies described in Fig. 1. For Phase II, in addition to the changes to the algorithm represented in SEEC-nt (see *Methods* section on “SEEC-nucleotide Algorithm”), we also optimized the selection temperature. Based on the rationale given in the text, we chose T = 1.5 and T = 0.75 for the SEEC-nt simulations using mfDCA and bmDCA parameters, respectively. All variant sequences are provided in Supplementary Tables 1 and 2.

### Variant Synthesis

Selected TEM-1 variants were cloned into a pET28a expression vector with inducible IPTG-controlled expression by the *lac* operon, along with an N-terminal His-tag and a kanamycin resistance gene. These expression vectors with our variant genes of interest were then each transformed into *E. coli* (BL21(DE3)) host cells. Gene synthesis of the TEM-1 sequence, mutagenesis of variants and plasmid cloning were performed by GenScript.

### Variant Strain *E. coli* Growth Assays

Individual transformed *E. coli* variant strains were grown in triplicates of cuvettes with 2 mL of culture volume to compare the rate of cell growth based on the change in optical density (OD) at 600nm over time. To begin the assay, cuvette setup involved adding 2 mL of Luria broth media, kanamycin (final concentration of 30 *μ*g/mL), IPTG (final concentration of 1mM), and either MIC of ampicillin (final concentration of 5 *μ*g/mL), a standard concentration (50 *μ*g/mL), or a challenging concentration (100 *μ*g/mL), depending on the set of variants being tested. Positive control sets of triplicate cuvettes were grown for each strain including WT, containing all the same culture reagents except for ampicillin. The negative control for every assay was an empty vector pET strain that contained all the same plasmid elements except for an ampicillin resistance gene. This empty vector strain was grown in triplicate containing all of the same reagents including equivalent levels of ampicillin. After culture setup, individual cuvettes were inoculated with 100 uL of overnight culture of each variant strain. Immediately after inoculation, culture OD was measured with a spectrophotometer, and readings were continued approximately every hour for a period of 24-72 hours. Final samples were collected and Sanger sequenced.

### Analysis of Growth Assay Data

Culture optical density (OD) at 600 nm was measured periodically for each strain over the period of 1-3 days to monitor cell growth and survival in the presence of ampicillin. The growth curve data points are the mean of 3 experimental replicates and error bars represent standard deviations. The maximum absorbance measurement of our spectrophotometer was 2. To analyze the fold growth during the growth assays, rationalized growth (*RG*) was calculated according to the equation:

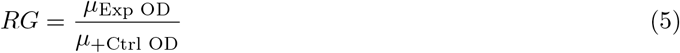

Where *μ*_Exp OD_ is the mean OD of the triplicate experimental cuvettes and *μ*_+Ctrl OD_ is the mean OD of the triplicate positive control cuvettes for each variant. The rationalized growth values for the variants were then normalized to the rationalized growth values of the WT strain to indicate the fold difference of growth (or Fold Growth, 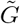 according to the equation:

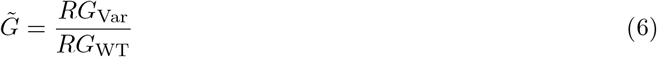

Where *RG*_Var_ is the rationalized growth for each variant and *RG*_WT_ is the rationalized growth of the WT strain. Fold difference in growth can be visualized with positive peaks illustrating enhanced growth and fold growth less than 1 represented sub-optimal function when compared to WT.

### Sanger Sequencing

After each growth assay, replicate sets of samples were combined and the cells pelleted by centrifugation. Plasmids were extracted via QIAprep spin miniprep kit (Qiagen) and quantitated using absorbance at 260 nm using a Nanodrop spectrophotometer. Relevant samples were then sent for Sanger sequencing performed by the Genome Center at The University of Texas at Dallas (Richardson, TX).

## Supporting information

Sanger Sequencing Chromatograms for Figure 3 data

Sanger Sequencing Chromatograms for Figure 4 data

Input Data and Raw Trajectory data

## Data Availability

All data used is publicly available. HMM seeds are available through PFAM [49]. Sequence databases used during *hmmsearch* are Swiss-Prot and TrEMBL from UniProt [52]. Training and input data available in Supplementary information.

## Code Availability

Code used in this work is available on Github at https://github.com/morcoslab/SEEC-NT.

## Acknowledgements

This research was funded by the University of Texas at Dallas (FM) the National Science Foundation CAREER grant number MCB-1943442 (FM), and the National Institutes of Health R35GM133631 (for FM, CMN and SA).

## Author Contributions

All authors conceived the study; C.M.N., N.M., T. H. and S.A. produced epistatic constraints; S.A. and C.M.N. performed experiments, produced and analyzed data. A.d.l.P. implemented the SEEC algorithms; S.A., C.M.N, A.d.l.P. and F.M. wrote the manuscript; FM supervised the work.

## References

[1] Jose Alberto de la Paz, Charisse M Nartey, Monisha Yuvaraj, and Faruck Morcos. Epistatic contributions promote the unification of incompatible models of neutral molecular evolution. Proceedings of the National Academy of Sciences, 117(11):5873–5882, 2020.

[2] Faruck Morcos, Andrea Pagnani, Bryan Lunt, Arianna Bertolino, Debora S Marks, Chris Sander, Riccardo Zecchina, José N Onuchic, Terence Hwa, and Martin Weigt. Direct-coupling analysis of residue coevolution captures native contacts across many protein families. Proceedings of the National Academy of Sciences, 108(49):E1293–E1301, 2011.

[3] Martin Weigt, Robert A White, Hendrik Szurmant, James A Hoch, and Terence Hwa. Identification of direct residue contacts in protein–protein interaction by message passing. Proceedings of the National Academy of Sciences, 106(1):67–72, 2009.

[4] Jiming Sheng and Shenshen Wang. Coevolutionary transitions emerging from flexible molecular recognition and eco-evolutionary feedback. Iscience, 24(8):102861, 2021.

[5] Muhammad Saqib Sohail, Raymond HY Louie, Zhenchen Hong, John P Barton, and Matthew R McKay. Inferring epistasis from genetic time-series data. Molecular Biology and Evolution, 39(10):msac199, 2022.

[6] David Ding, Anna G Green, Boyuan Wang, Thuy-Lan Vo Lite, Eli N Weinstein, Debora S Marks, and Michael T Laub. Co-evolution of interacting proteins through non-contacting and non-specific mutations. Nature ecology & evolution, 6(5):590–603, 2022.

[7] Joseph L Harman, Patrick N Reardon, Shawn M Costello, Gus D Warren, Sophia R Phillips, Patrick J Connor, Susan Marqusee, and Michael J Harms. Evolution avoids a pathological stabilizing interaction in the immune protein s100a9. Proceedings of the National Academy of Sciences, 119(41):e2208029119, 2022.

[8] Joseph D Bryngelson and Peter G Wolynes. Spin glasses and the statistical mechanics of protein folding. Proceedings of the National Academy of sciences, 84(21):7524–7528, 1987.

[9] José Nelson Onuchic, Zaida Luthey-Schulten, and Peter G Wolynes. Theory of protein folding: the energy landscape perspective. Annual review of physical chemistry, 48(1):545–600, 1997.

[10] Alan Lapedes, Bertrand Giraud, and Christopher Jarzynski. Using sequence alignments to predict protein structure and stability with high accuracy. LANL preprint LA-UR-02-4481, 2002.

[11] Omar Haq, Michael Andrec, Alexandre V Morozov, and Ronald M Levy. Correlated electrostatic mutations provide a reservoir of stability in hiv protease. 2012.

[12] Faruck Morcos, Nicholas P Schafer, Ryan R Cheng, José N Onuchic, and Peter G Wolynes. Coevolutionary information, protein folding landscapes, and the thermodynamics of natural selection. Proceedings of the National Academy of Sciences, 111(34):12408–12413, 2014.

[13] Ronald M Levy, Allan Haldane, and William F Flynn. Potts hamiltonian models of protein co-variation, free energy landscapes, and evolutionary fitness. Current opinion in structural biology, 43:55–62, 2017.

[14] Thomas A Hopf, John B Ingraham, Frank J Poelwijk, Charlotta PI Schärfe, Michael Springer, Chris Sander, and Debora S Marks. Mutation effects predicted from sequence co-variation. Nature biotechnology, 35(2):128–135, 2017.

[15] Matteo Figliuzzi, Hervé Jacquier, Alexander Schug, Oliver Tenaillon, and Martin Weigt. Coevolutionary landscape inference and the context-dependence of mutations in beta-lactamase tem-1. Molecular biology and evolution, 33(1):268–280, 2016.

[16] Xian-Li Jiang, Rey P Dimas, Clement TY Chan, and Faruck Morcos. Coevolutionary methods enable robust design of modular repressors by reestablishing intra-protein interactions. Nature communications, 12(1):5592, 2021.

[17] Arvind Murugan, Kabir Husain, Michael J Rust, Chelsea Hepler, Joseph Bass, Julian MJ Pietsch, Peter S Swain, Siddhartha G Jena, Jared E Toettcher, Arup K Chakraborty, et al. Roadmap on biology in time varying environments. Physical biology, 18(4):041502, 2021.

[18] Michael A Stiffler, Frank J Poelwijk, Kelly P Brock, Richard R Stein, Adam Riesselman, Joan Teyra, Sachdev S Sidhu, Debora S Marks, Nicholas P Gauthier, and Chris Sander. Protein structure from experimental evolution. Cell Systems, 10(1):15–24, 2020.

[19] Marco Fantini, Simonetta Lisi, Paolo De Los Rios, Antonino Cattaneo, and Annalisa Pastore. Protein structural information and evolutionary landscape by in vitro evolution. Molecular biology and evolution, 37(4):1179–1192, 2020.

[20] Matteo Figliuzzi, Pierre Barrat-Charlaix, and Martin Weigt. How pairwise coevolutionary models capture the collective residue variability in proteins? Molecular biology and evolution, 35(4):1018–1027, 2018.

[21] William P Russ, Matteo Figliuzzi, Christian Stocker, Pierre Barrat-Charlaix, Michael Socolich, Peter Kast, Donald Hilvert, Remi Monasson, Simona Cocco, Martin Weigt, et al. An evolution-based model for designing chorismate mutase enzymes. Science, 369(6502):440–445, 2020.

[22] Pengfei Tian, Adrien Lemaire, Fabien Sénéchal, Olivier Habrylo, Viviane Antonietti, Pascal Sonnet, Valérie Lefebvre, Frederikke Isa Marin, Robert B Best, Jérôme Pelloux, et al. Design of a protein with improved thermal stability by an evolution-based generative model. Angewandte Chemie International Edition, page e202202711, 2022.

[23] Cheyenne Ziegler, Jonathan Martin, Claude Sinner, and Faruck Morcos. Latent generative landscapes as maps of functional diversity in protein sequence space. Nature Communications, 14(1):2222, 2023.

[24] Matteo Bisardi, Juan Rodriguez-Rivas, Francesco Zamponi, and Martin Weigt. Modeling sequence-space exploration and emergence of epistatic signals in protein evolution. Molecular biology and evolution, 39(1):msab321, 2022.

[25] Caroline M Weisman. The origins and functions of de novo genes: Against all odds? Journal of Molecular Evolution, 90(3-4):244–257, 2022.

[26] Diane Marie Keeling, Patricia Garza, Charisse Michelle Nartey, and Anne-Ruxandra Carvunis. The meanings of’function’in biology and the problematic case of de novo gene emergence. Elife, 8:e47014, 2019.

[27] Lucile Vigué, Giancarlo Croce, Marie Petitjean, Etienne Ruppé, Olivier Tenaillon, and Martin Weigt. Deciphering polymorphism in 61,157 escherichia coli genomes via epistatic sequence landscapes. Nature Communications, 13(1):4030, 2022.

[28] Stephen F Altschul, Warren Gish, Webb Miller, Eugene W Myers, and David J Lipman. Basic local alignment search tool. Journal of molecular biology, 215(3):403–410, 1990.

[29] Nobuhiko Tokuriki and Dan S Tawfik. Stability effects of mutations and protein evolvability. Current opinion in structural biology, 19(5):596–604, 2009.

[30] Shimon Bershtein, Michal Segal, Roy Bekerman, Nobuhiko Tokuriki, and Dan S Tawfik. Robustness–epistasis link shapes the fitness landscape of a randomly drifting protein. Nature, 444(7121):929–932, 2006.

[31] Shamil Sunyaev, Vasily Ramensky, Ina Koch, Warren Lathe III, Alexey S Kondrashov, and Peer Bork. Prediction of deleterious human alleles. Human molecular genetics, 10(6):591–597, 2001.

[32] Rachel Karchin, Mark Diekhans, Libusha Kelly, Daryl J Thomas, Ursula Pieper, Narayanan Eswar, David Haussler, and Andrej Sali. Ls-snp: large-scale annotation of coding non-synonymous snps based on multiple information sources. Bioinformatics, 21(12):2814–2820, 2005.

[33] Yana Bromberg and Burkhard Rost. Snap: predict effect of non-synonymous polymorphisms on function. Nucleic acids research, 35(11):3823–3835, 2007.

[34] Natalie CJ Strynadka, Susan E Jensen, Kathy Johns, Helen Blanchard, Malcolm Page, André Matagne, Jean-Marie Frere, and Michael NG James. Structural and kinetic characterization of a β-lactamase-inhibitor protein. Nature, 368(6472):657–660, 1994.

[35] Natalie CJ Strynadka, Susan E Jensen, Pedro M Alzari, and Michael NG James. A potent new mode of β-lactamase inhibition revealed by the 1.7 a x-ray crystallographic structure of the tem-1–blip complex. Nature structural biology, 3(3):290–297, 1996.

[36] Bartlomiej G Fryszczyn, Carolyn J Adamski, Nicholas G Brown, Kacie Rice, Wanzhi Huang, and Timothy Palzkill. Role of β-lactamase residues in a common interface for binding the structurally unrelated inhibitory proteins blip and blip-ii. Protein Science, 23(9):1235–1246, 2014.

[37] Thomas D Townsley, James T Wilson, Harrison Akers, Timothy Bryant, Salvador Cordova, TL Wallace, Kirk K Durston, and Joseph E Deweese. Psicalc: a novel approach to identifying and ranking critical nonproximal interdependencies within the overall protein structure. Bioinformatics Advances, 2(1):vbac058, 2022.

[38] Najeeb Halabi, Olivier Rivoire, Stanislas Leibler, and Rama Ranganathan. Protein sectors: evolutionary units of three-dimensional structure. Cell, 138(4):774–786, 2009.

[39] Michael Schmidt and Kay Hamacher. hodca: higher order direct-coupling analysis. BMC bioinformatics, 19(1):1–5, 2018.

[40] Yusuf Talha Tamer, Ilona K Gaszek, Haleh Abdizadeh, Tugce Altinusak Batur, Kimberly A Reynolds, Ali Rana Atilgan, Canan Atilgan, and Erdal Toprak. High-order epistasis in catalytic power of dihydrofolate reductase gives rise to a rugged fitness landscape in the presence of trimethoprim selection. Molecular biology and evolution, 36(7):1533–1550, 2019.

[41] Paul E O’maille, Arthur Malone, Nikki Dellas, B Andes Hess Jr, Lidia Smentek, Iseult Sheehan, Bryan T Greenhagen, Joe Chappell, Gerard Manning, and Joseph P Noel. Quantitative exploration of the catalytic landscape separating divergent plant sesquiterpene synthases. Nature chemical biology, 4(10):617–623, 2008.

[42] Aditya Ballal, Caroline Laurendon, Melissa Salmon, Maria Vardakou, Jitender Cheema, Marianne Defernez, Paul E O’Maille, and Alexandre V Morozov. Sparse epistatic patterns in the evolution of terpene synthases. Molecular biology and evolution, 37(7):1907–1924, 2020.

[43] Gary W Rudgers and Timothy Palzkill. Identification of residues in β-lactamase critical for binding β-lactamase inhibitory protein. Journal of Biological Chemistry, 274(11):6963–6971, 1999.

[44] Raymond HY Louie, Kevin J Kaczorowski, John P Barton, Arup K Chakraborty, and Matthew R McKay. Fitness landscape of the human immunodeficiency virus envelope protein that is targeted by antibodies. Proceedings of the National Academy of Sciences, 115(4):E564–E573, 2018.

[45] Tian-hao Zhang, Lei Dai, John P Barton, Yushen Du, Yuxiang Tan, Wenwen Pang, Arup K Chakraborty, James O Lloyd-Smith, and Ren Sun. Predominance of positive epistasis among drug resistance-associated mutations in hiv-1 protease. PLoS genetics, 16(10):e1009009, 2020.

[46] Hong-Li Zeng, Vito Dichio, Edwin Rodríguez Horta, Kaisa Thorell, and Erik Aurell. Global analysis of more than 50,000 sars-cov-2 genomes reveals epistasis between eight viral genes. Proceedings of the National Academy of Sciences, 117(49):31519–31526, 2020.

[47] Juan Rodriguez-Rivas, Giancarlo Croce, Maureen Muscat, and Martin Weigt. Epistatic models predict mutable sites in sars-cov-2 proteins and epitopes. Proceedings of the National Academy of Sciences, 119(4):e2113118119, 2022.

[48] Joao V Rodrigues and Eugene I Shakhnovich. Adaptation to mutational inactivation of an essential gene converges to an accessible suboptimal fitness peak. Elife, 8:e50509, 2019.

[49] Robert D Finn, Alex Bateman, Jody Clements, Penelope Coggill, Ruth Y Eberhardt, Sean R Eddy, Andreas Heger, Kirstie Hetherington, Liisa Holm, Jaina Mistry, et al. Pfam: the protein families database. Nucleic acids research, 42(D1):D222–D230, 2014.

[50] Sean R Eddy. Accelerated profile hmm searches. PLoS computational biology, 7(10):e1002195, 2011.

[51] Sean R. Eddy. HMMER: 3.3.2. http://hmmer.org, November 2020.

[52] Uniprot: the universal protein knowledgebase in 2021. Nucleic acids research, 49(D1):D480–D489, 2021.

[53] Ryan R Cheng, Olle Nordesjö, Ryan L Hayes, Herbert Levine, Samuel C Flores, José N Onuchic, and Faruck Morcos. Connecting the sequence-space of bacterial signaling proteins to phenotypes using coevolutionary landscapes. Molecular biology and evolution, 33(12):3054–3064, 2016.

[54] Qin Zhou, Nikesh Kunder, José Alberto De la Paz, Alexandra E Lasley, Vandita D Bhat, Faruck Morcos, and Zachary T Campbell. Global pairwise rna interaction landscapes reveal core features of protein recognition. Nature communications, 9(1):2511, 2018.

[55] Krithika Ravishankar, Xianli Jiang, Emmett M Leddin, Faruck Morcos, and G Andrés Cisneros. Computational compensatory mutation discovery approach: Predicting a parp1 variant rescue mutation. Biophysical Journal, 121(19):3663–3673, 2022.

