## Supplementary information for "*In vivo* functional phenotypes from a computational epistatic model of evolution"

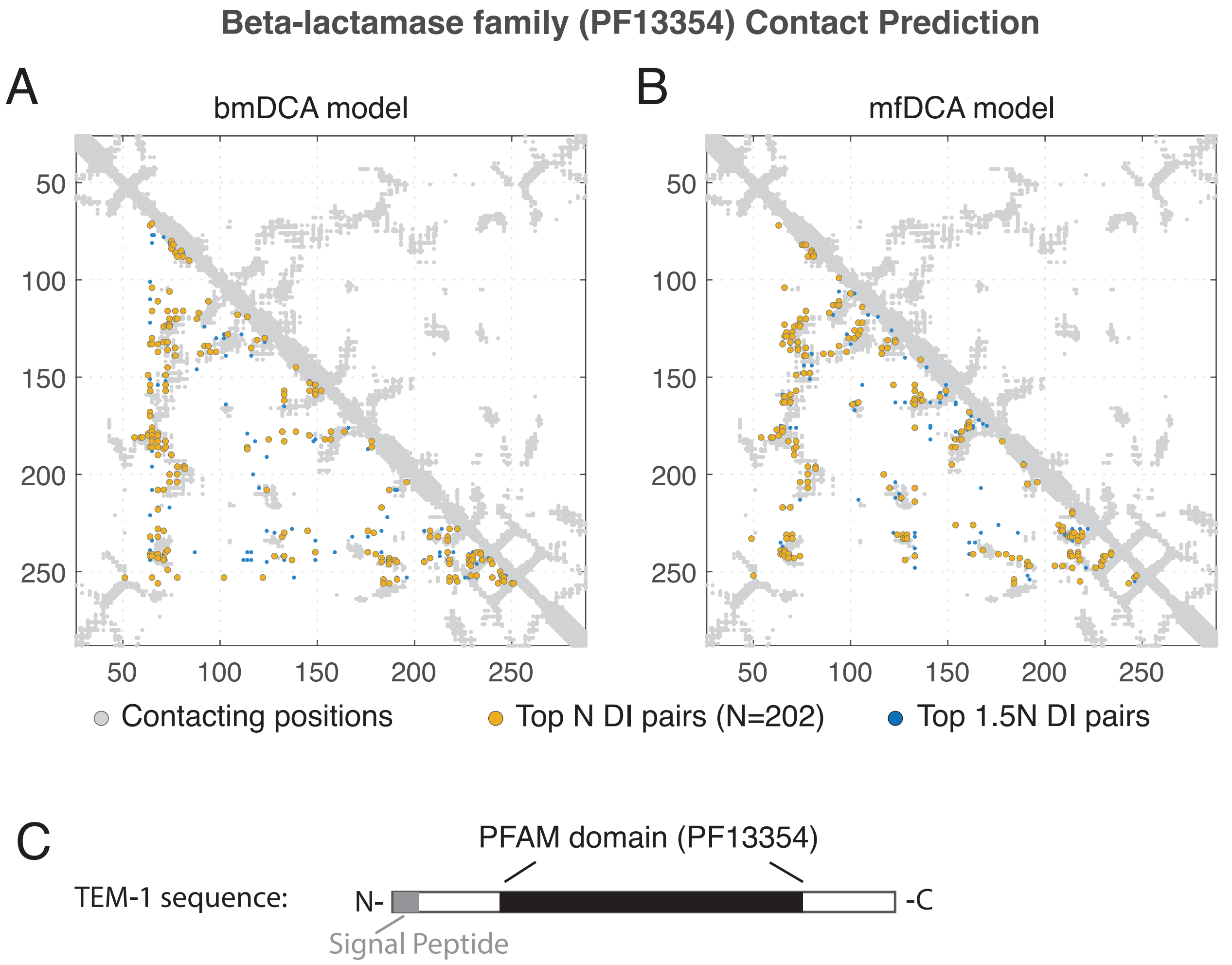


**Supplementary Figure 1:** Validation of mf (A) and bmDCA (B) models through prediction of structural contacts. The top N and 1.5N DI pairs capture structural contacts in the *E. coli* TEM-1 structure (PDB ID: 1ERM). N is the length of the model, which is defined by the PFAM domain PF13354(N=202). (C) Schematic of the model used for the MSA (black area) relative to the *E. coli* TEM-1 gene.

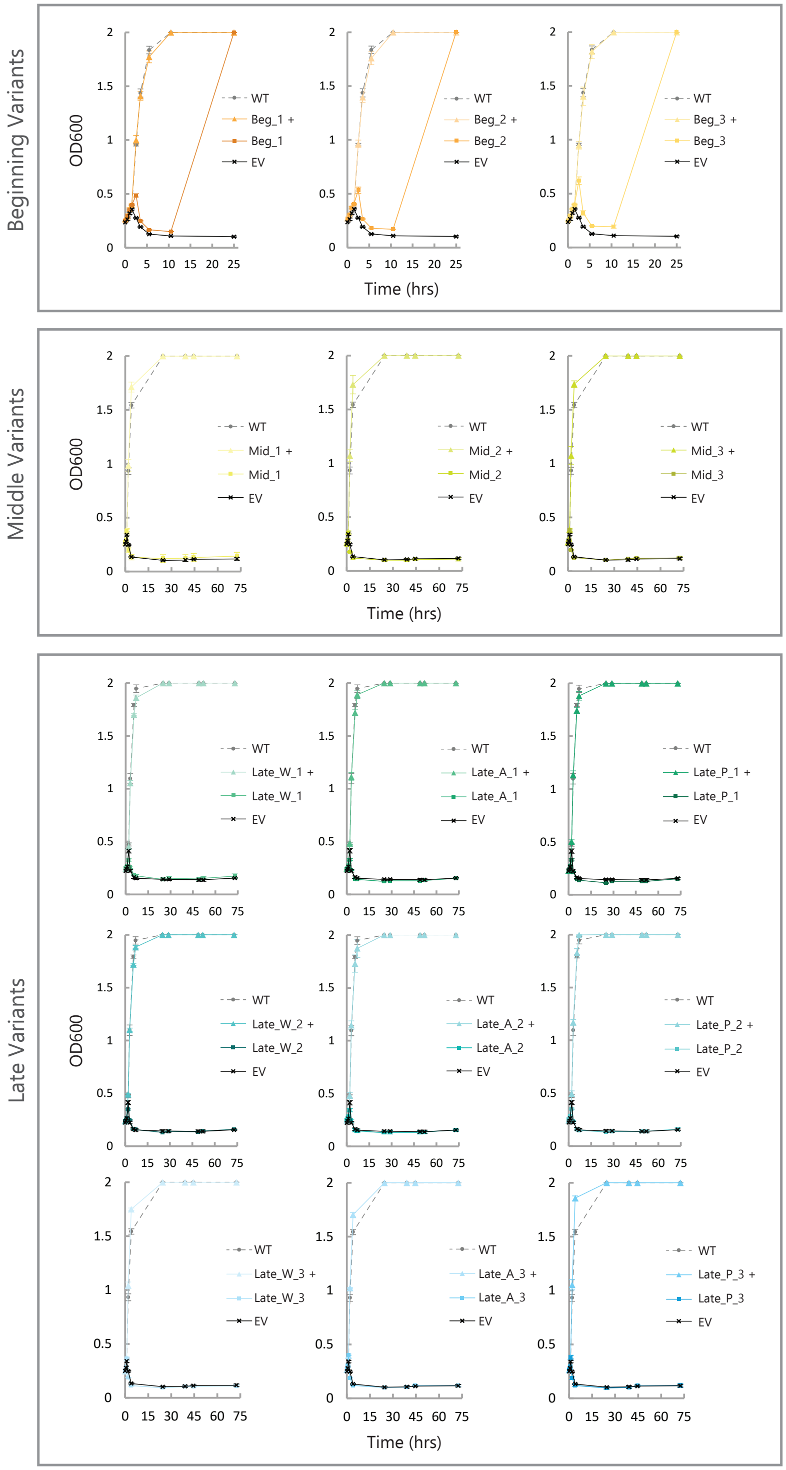

**Supplementary Figure 2:** Individual *E. coli* growth curves for variants picked from the **bmDCA** model SEEC-aa trajectory. Cultures were grown in 5 $\mu$g/mL Ampicillin. In the positive controls (+), variants were grown in the absence of Ampicillin. Data points are the mean of 3 experimental replicates and error bars represent standard deviations.

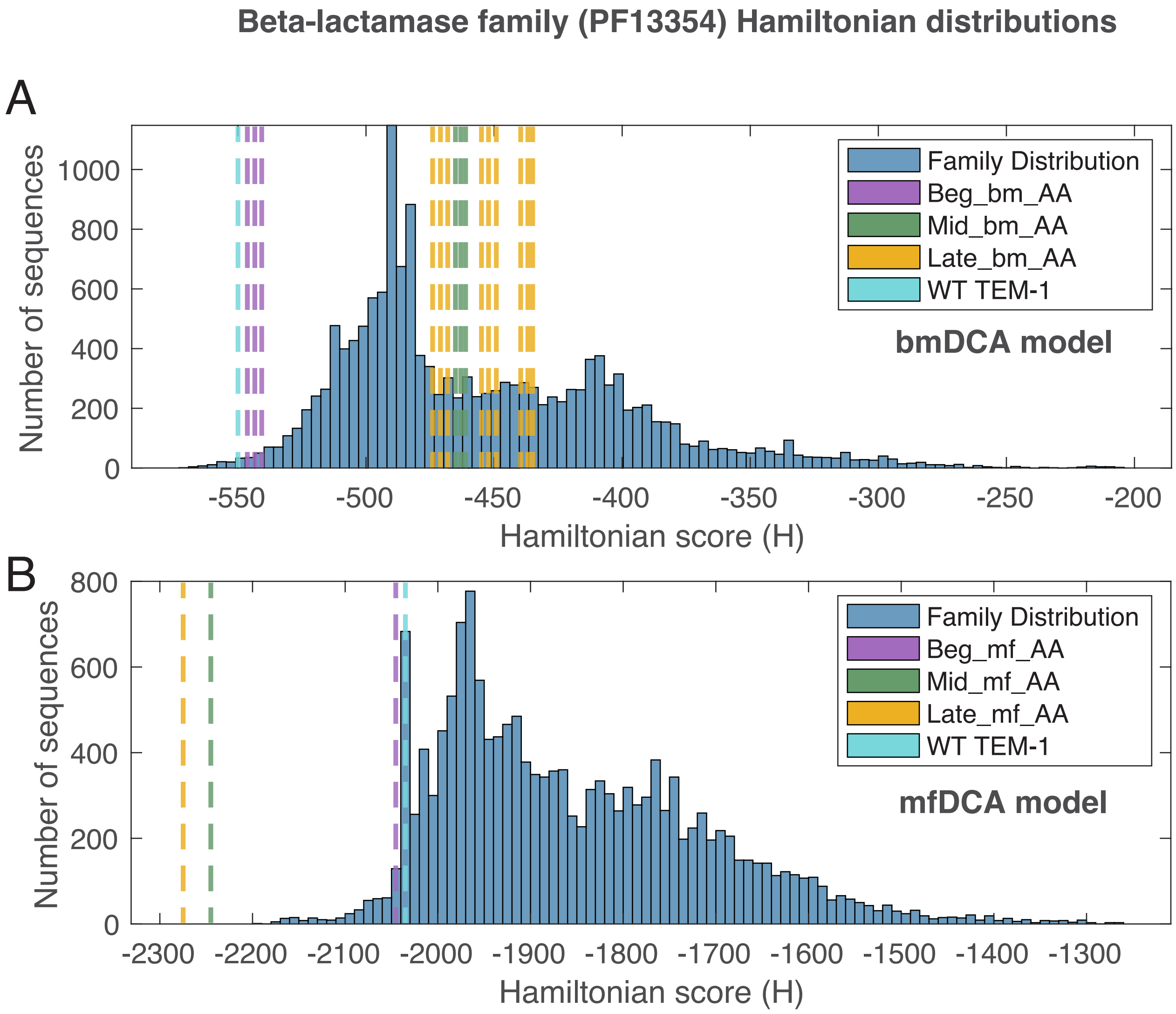


**Supplementary Figure 3: PFAM model Family Hamiltonian Distributions** Comparison of $H$ scores for Native family members relative to variants chosen from SEEC-aa simulations. The two panels are for the mean field DCA (A) and Boltzmann machine learning DCA (B) based models. Variants are divided into three groups, beginning, middle and late. The $H$ score of the WT TEM-1 sequence is shown for comparison. $H$ scores of Mid and Late mfDCA SEEC-aa variants are favorable, but outside the distribution of Native protein scores.

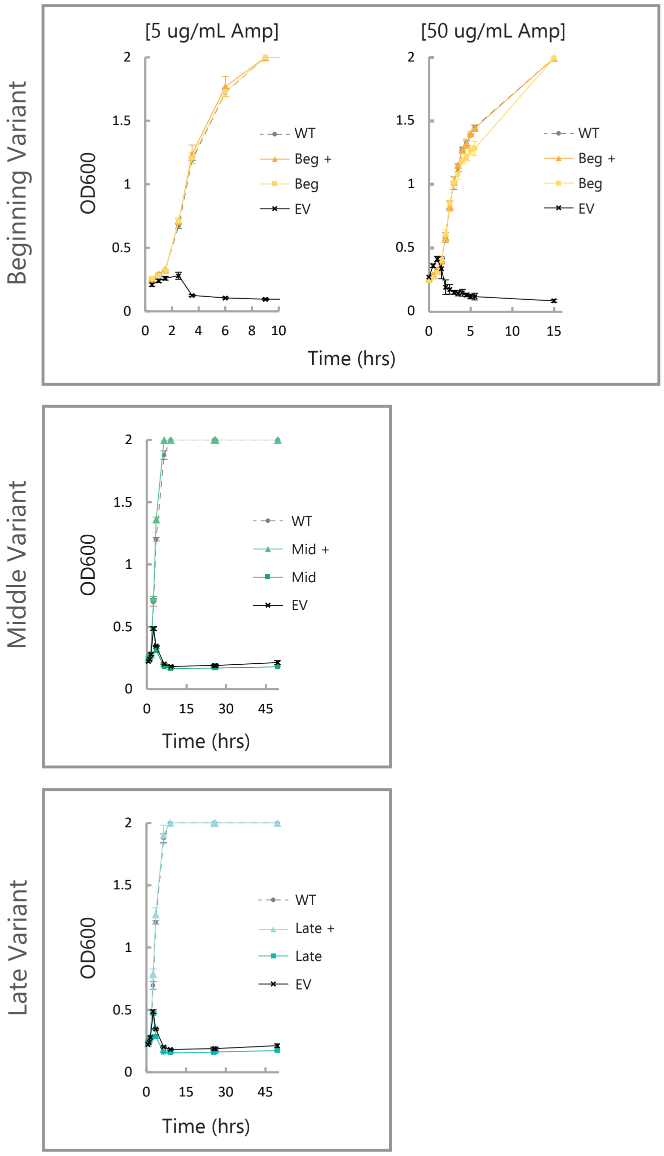


**Supplementary Figure 4:** Individual *E. coli* growth curves for variants picked from the **mfDCA** model SEEC-aa trajectory. Beginning variant cultures were grown in 5 and 50 $\mu$g/mL Ampicillin, middle and late variant cultures were grown in 5 $\mu$g/mL Ampicillin. In the positive controls (+), variants were grown in the absence of Ampicillin. Data points are the mean of 3 experimental replicates and error bars represent standard deviations.

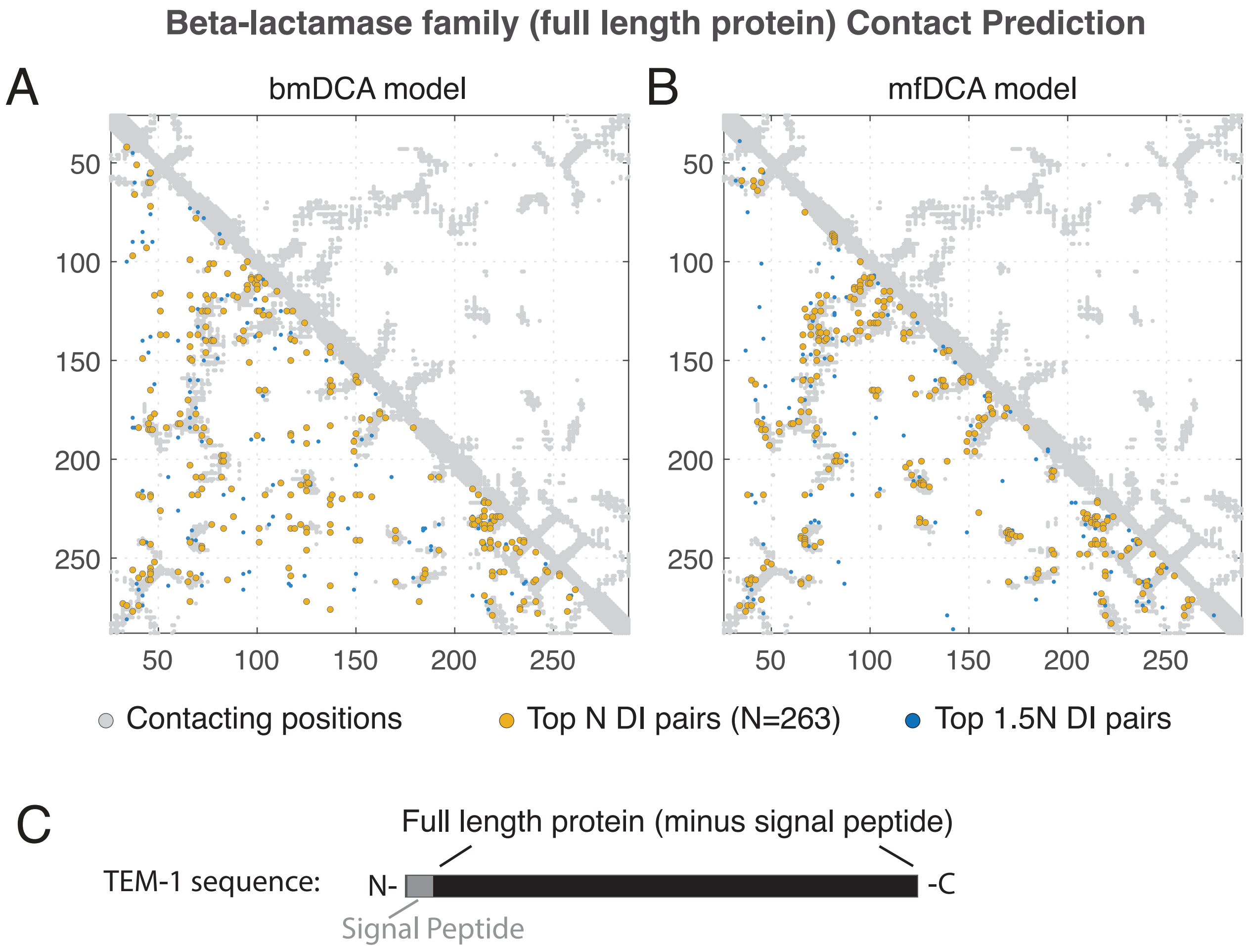


**Supplementary Figure 5:** Validation of mf (A) and bmDCA (B) models through prediction of structural contacts. The top N and 1.5N DI pairs capture structural contacts in the *E. coli* TEM-1 structure (PDB ID: 1ERM). N is the length of the model, which is the full length protein minus the signal peptide (N=263 residues). (C) Schematic of the model used for the MSA (black area) relative to the *E. coli* TEM-1 gene.

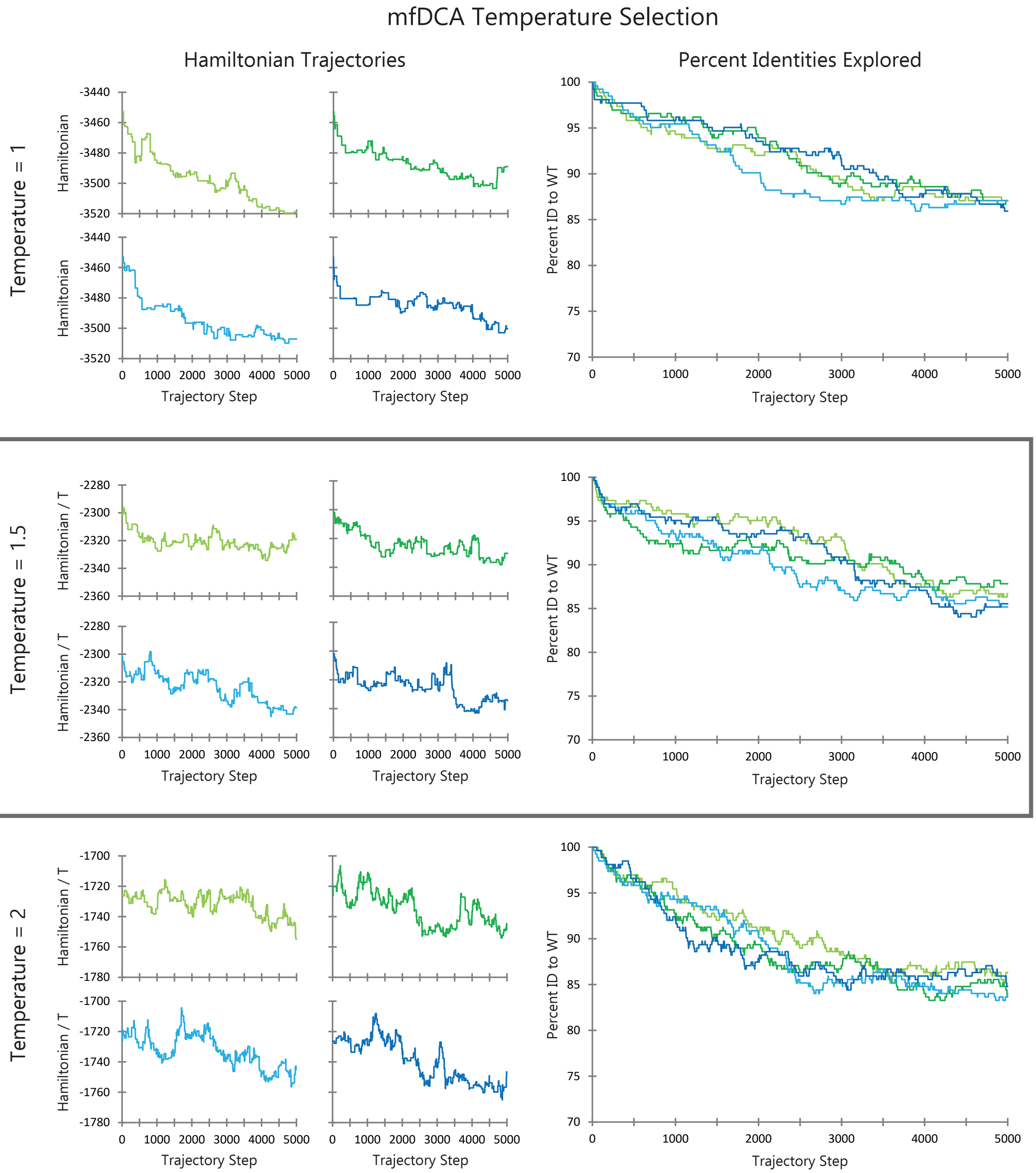


**Supplementary Figure 6:** Testing the effects of various temperatures for mfDCA informed SEEC-nt simulations. The number of steps for each simulation is 5000. The preferred temperature is 1.5 with the best allowance for sequence space exploration while still trending downwards in Hamiltonian relative to the simulation temperature (outlined in the gray box).

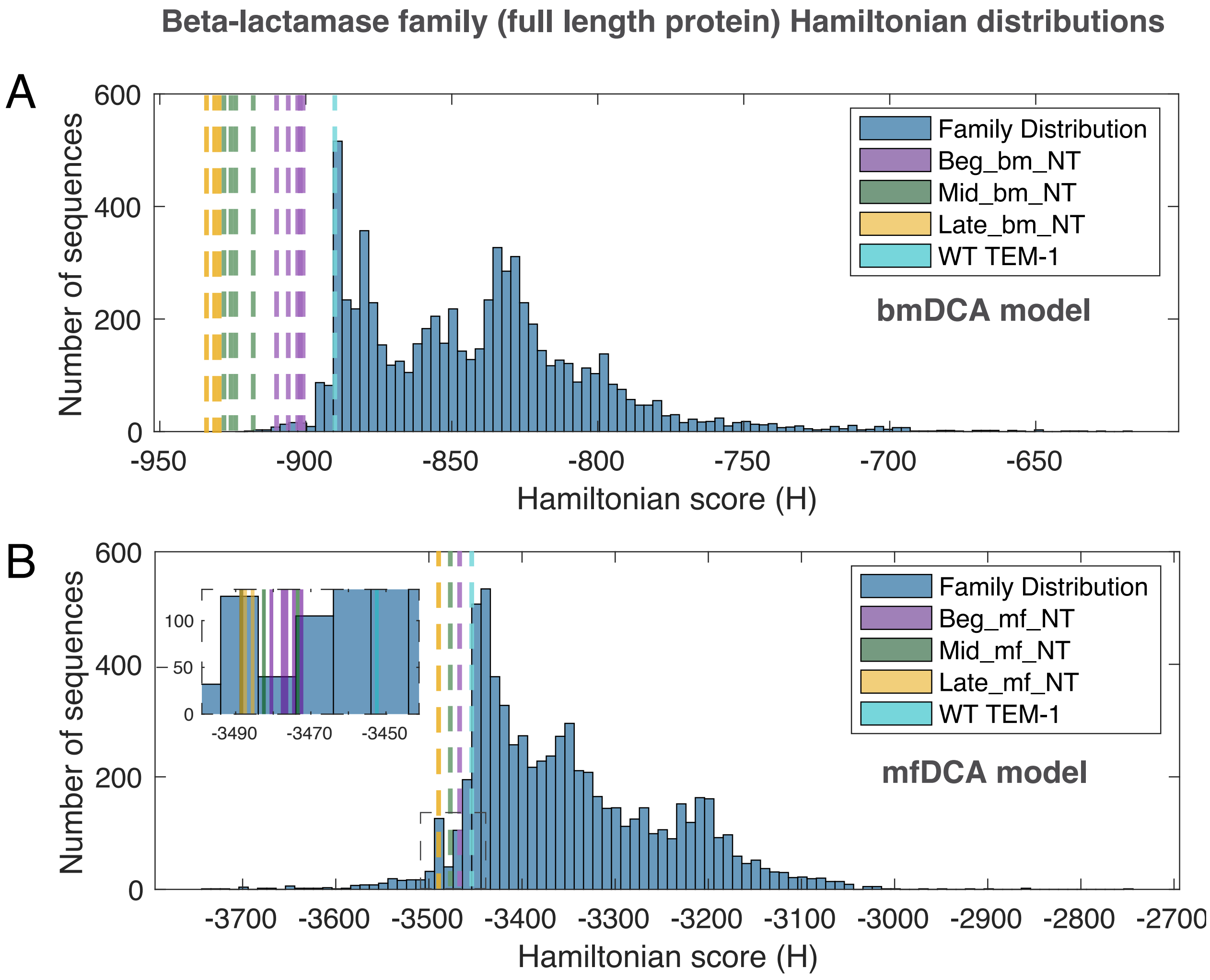


**Supplementary Figure 7: Full Length Protein Family Hamiltonian Distributions** Comparison of $H$ scores for Native family members relative to variants chosen from SEEC-nt simulations. Variants are divided into three groups, beginning, middle and late. The $H$ score of the WT TEM-1 sequence is shown for comparison. (A) Mean-field DCA based model. Each dotted line represents all the variants in the Beg, Mid or Late groups. The inset has one solid line per variant: 6 Beg, 4 Mid, 3 Late variants.(B) Boltzmann machine learning DCA based model. $H$ scores of Mid and Late mfDCA SEEC-nt variants are favorable, but outside the distribution of Native protein scores.

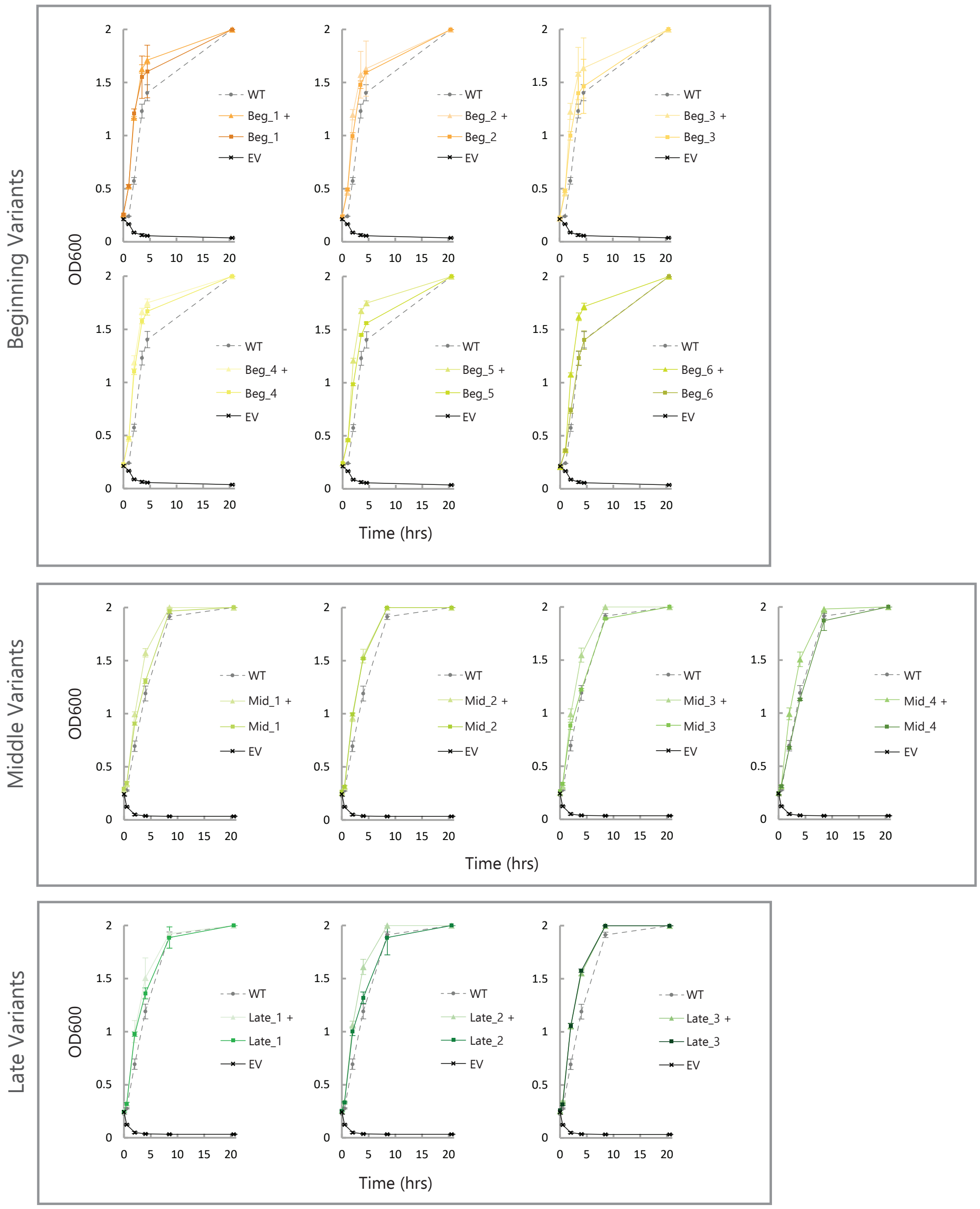


**Supplementary Figure 8:** Individual *E. coli* growth curves for variants picked from the **mfDCA** model SEEC-nt trajectory. Cultures were grown in 50 $\mu$g/mL Ampicillin. In the positive controls (+), variants were grown in the absence of Ampicillin. Data points are the mean of 3 experimental replicates and error bars represent standard deviations.

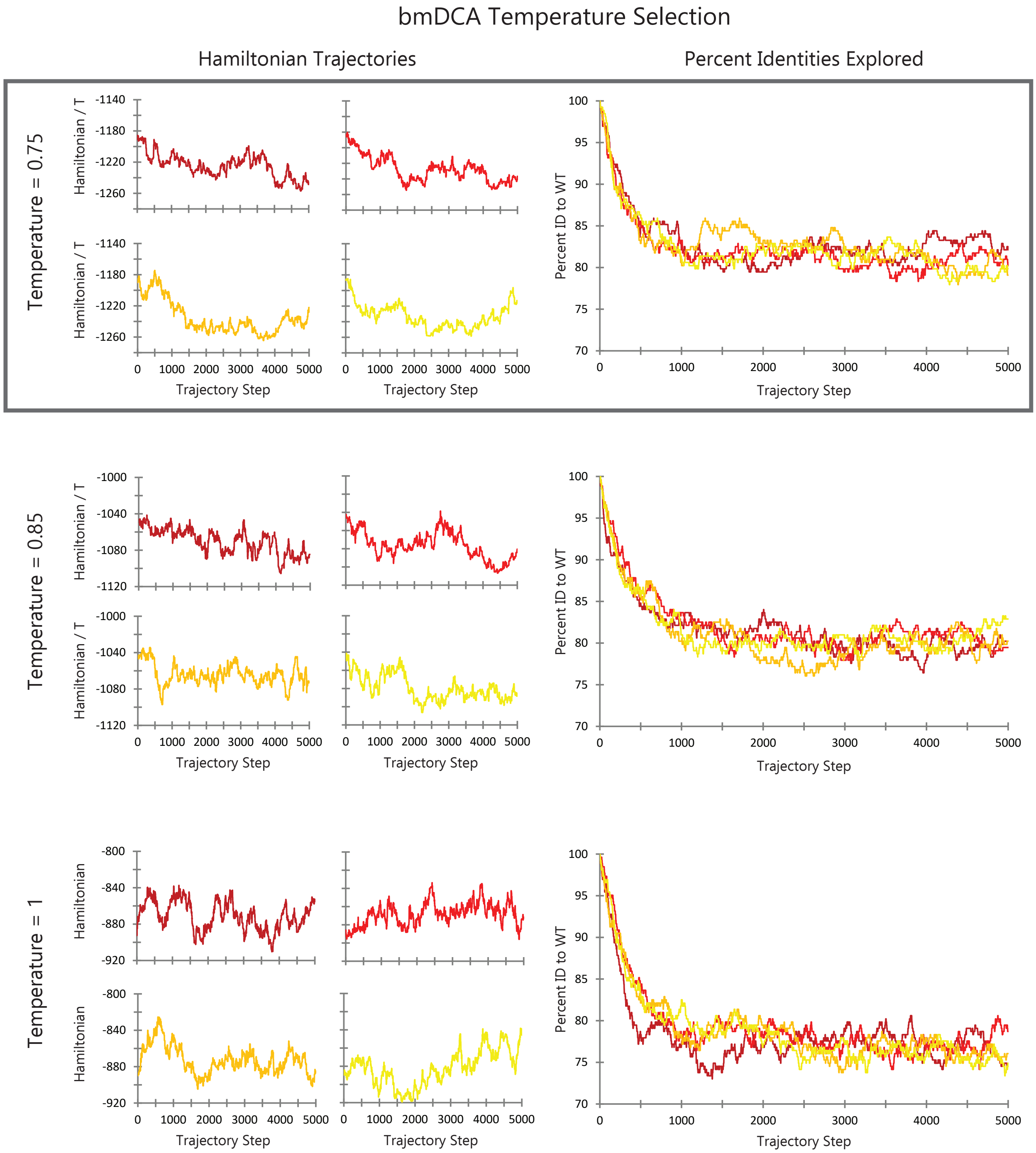


**Supplementary Figure 9:** Testing the effects of various temperatures for bmDCA informed SEEC-nt simulations. The number of steps for each simulation is 5000. The preferred temperature is 0.75 with the best allowance for sequence space exploration while still trending downwards in Hamiltonian relative to the simulation temperature (outlined in the gray box).

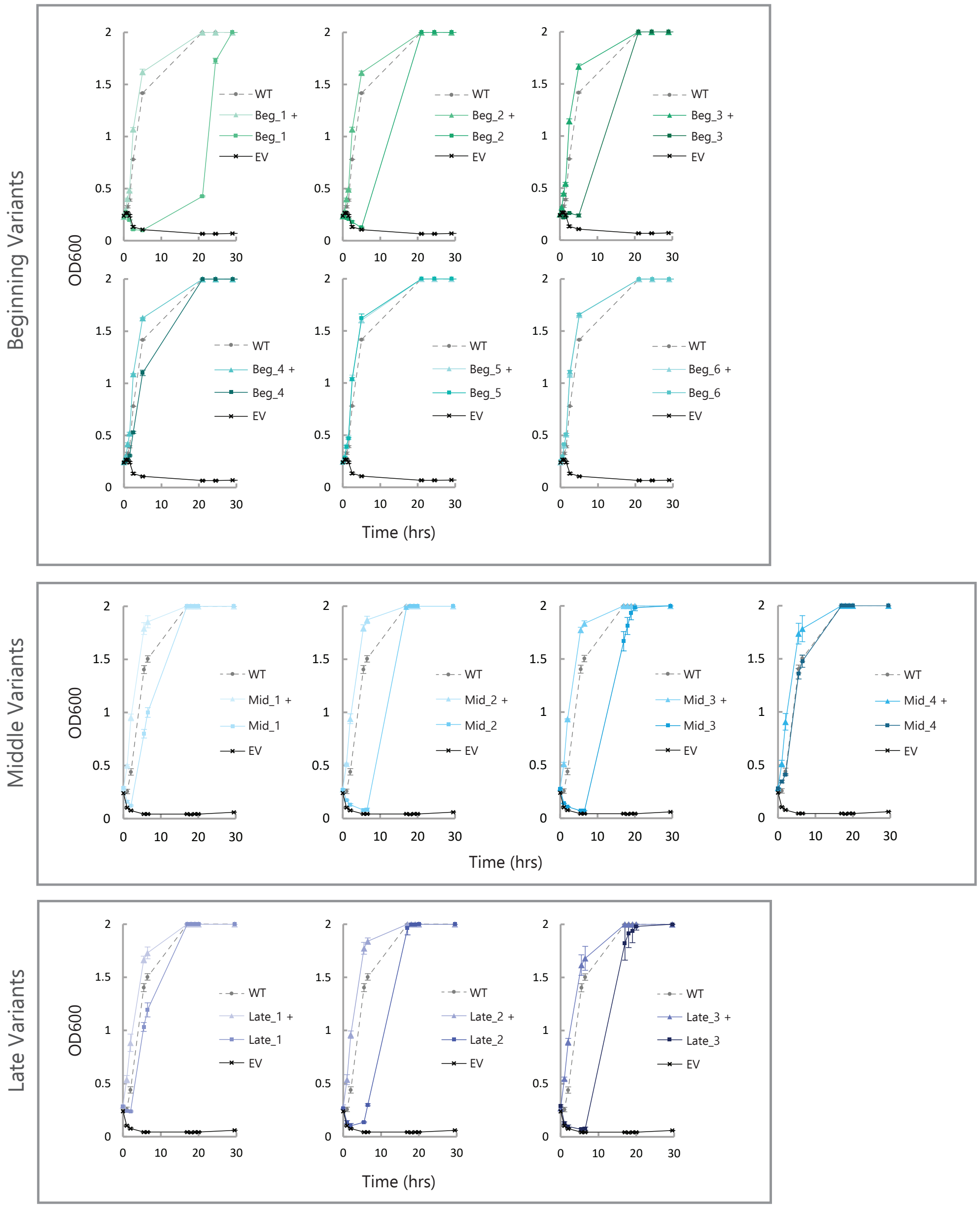


**Supplementary Figure 10:** Individual *E. coli* growth curves for variants picked from the **bmDCA** model SEEC-nt trajectory. Cultures were grown in 50 $\mu$g/mL Ampicillin. In the positive controls (+), variants were grown in the absence of Ampicillin. Data points are the mean of 3 experimental replicates and error bars represent standard deviations.

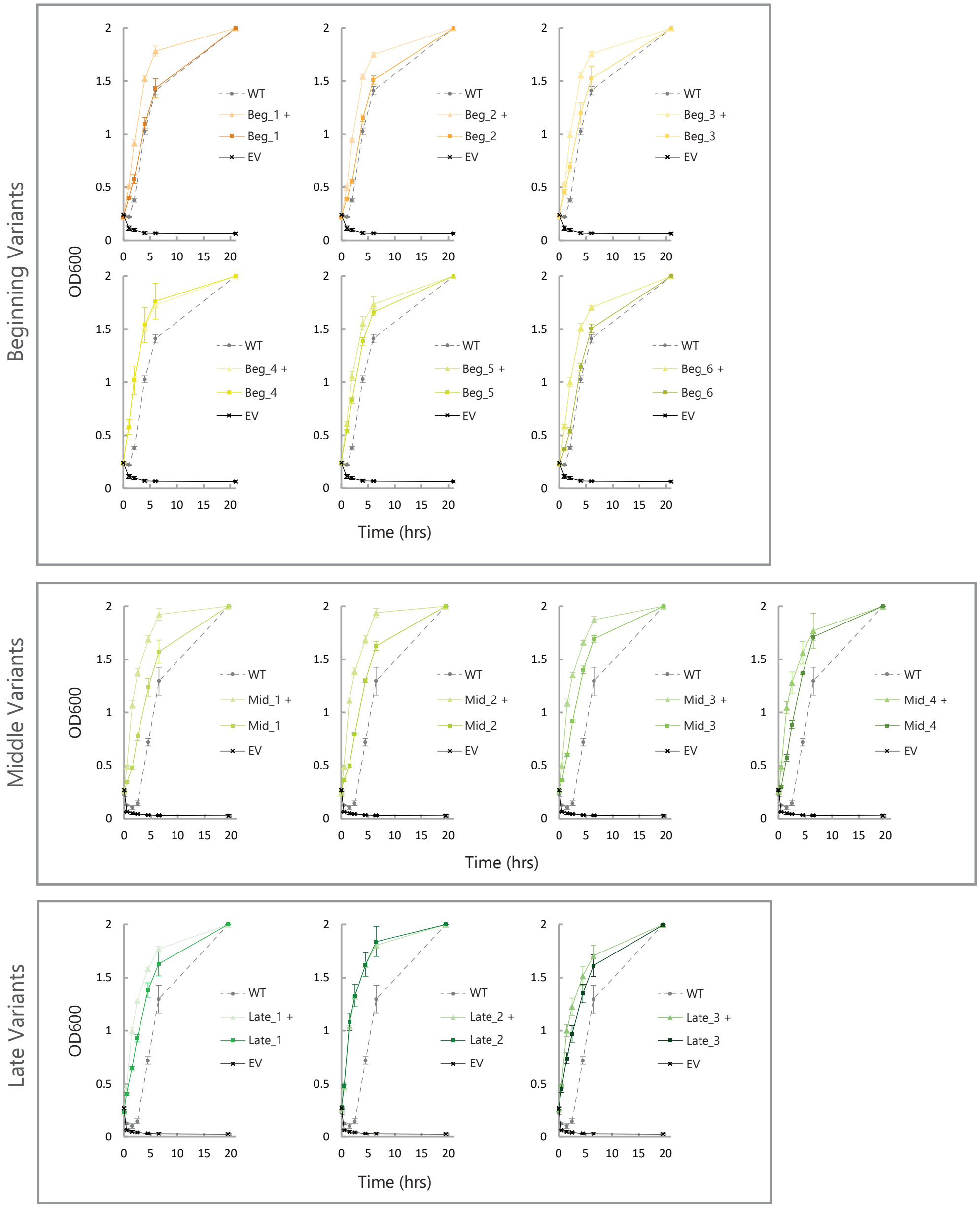


**Supplementary Figure 11:** Individual *E. coli* growth curves for variants picked from the **mfDCA** model SEEC-nt trajectory. Cultures were grown in 100 $\mu$g/mL Ampicillin. In the positive controls (+), variants were grown in the absence of Ampicillin. Data points are the mean of 3 experimental replicates and error bars represent standard deviations.

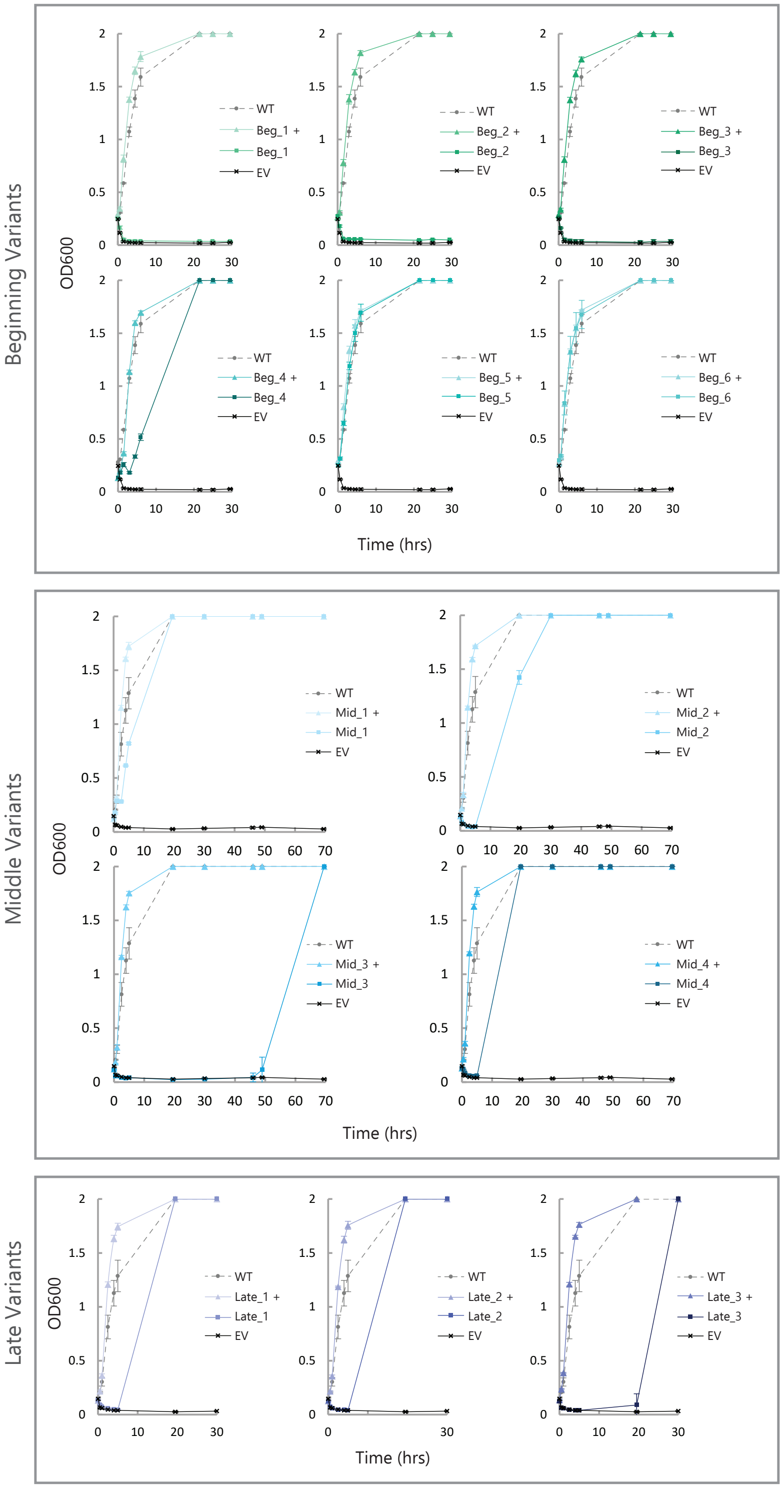


**Supplementary Figure 12:** Individual *E. coli* growth curves for variants picked from the **bmDCA** model SEEC-nt trajectory. Cultures were grown in 100 $\mu$g/mL Ampicillin. In the positive controls (+), variants were grown in the absence of Ampicillin. Data points are the mean of 3 experimental replicates and error bars represent standard deviations.

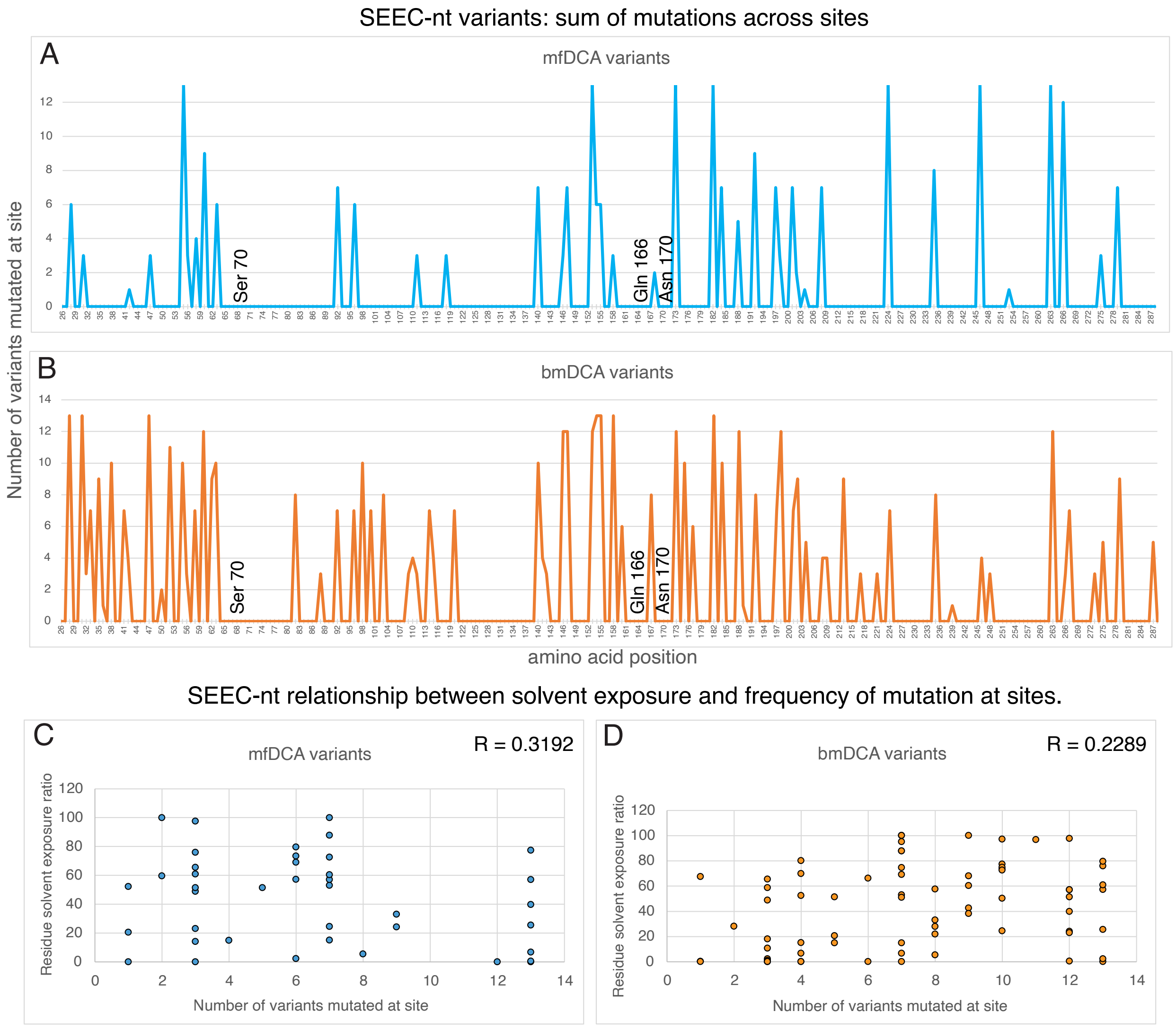


**Supplementary Figure 13:** Comparison of sequence space exploration within mean field (A) and Boltzmann machine learning (B) variants. Mutations per site are plotted and numbering assumes the first residue after the signal peptide is position 1. Conserved catalytic residues are labeled.(C,D)Relationship between Ratio of solvent exposure for each residue vs the number of times it was mutated among the variants for mfDCA(C) and bmDCA(D) variants. Solvent exposed surface area ratios calculated using ref. [1] with the default settings.

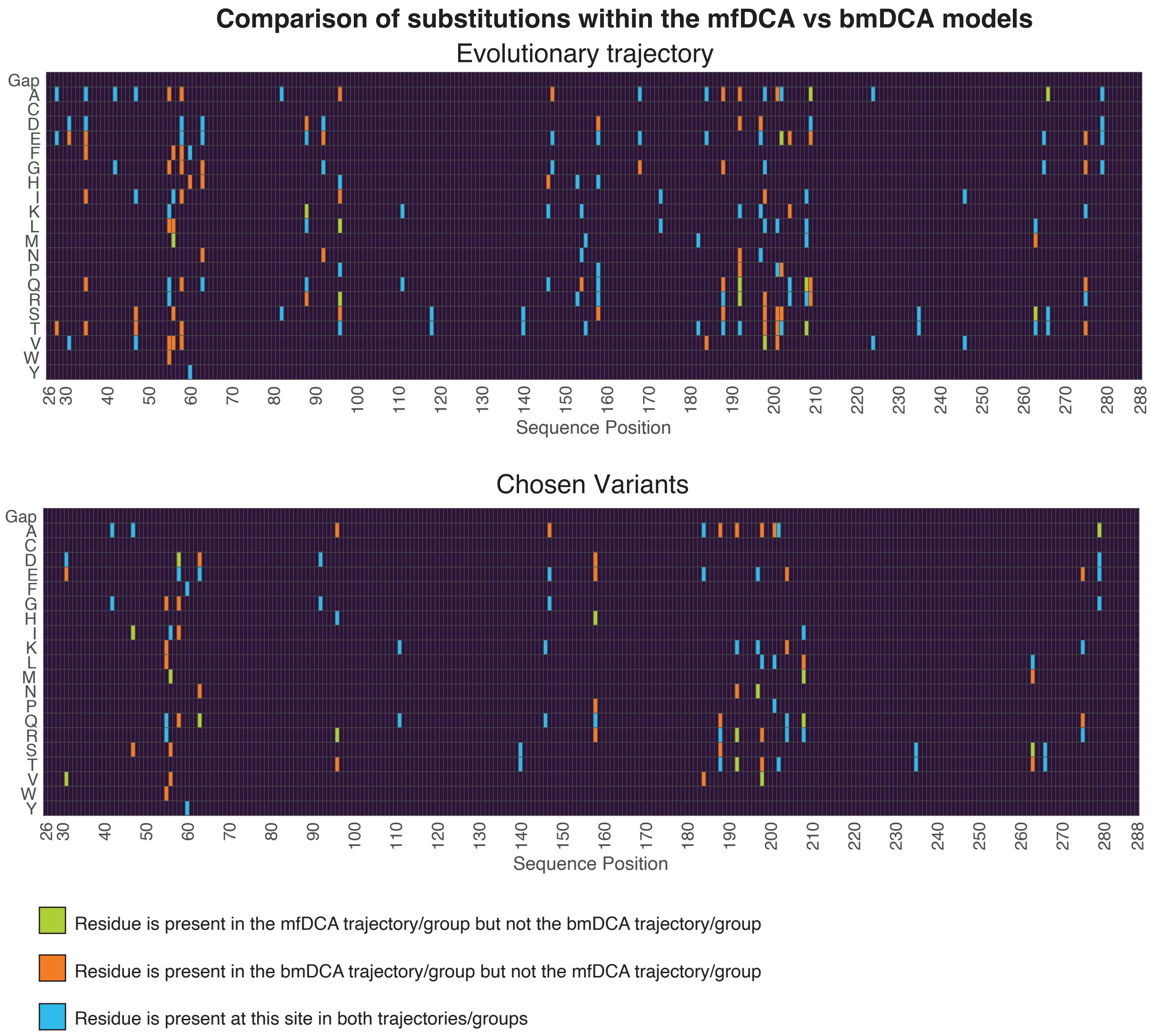


**Supplementary Figure 14:** Comparisons of substitutions with the mfDCA and bmDCA models for sequences generated by the SEEC-nt trajectory or the subset chosen for experimental testing. Columns with only black rectangles were conserved in either the *bm*, *mf* model or both; this plot focuses on which particular residues are unique to one model or shared by both when a substitution is made at a site.

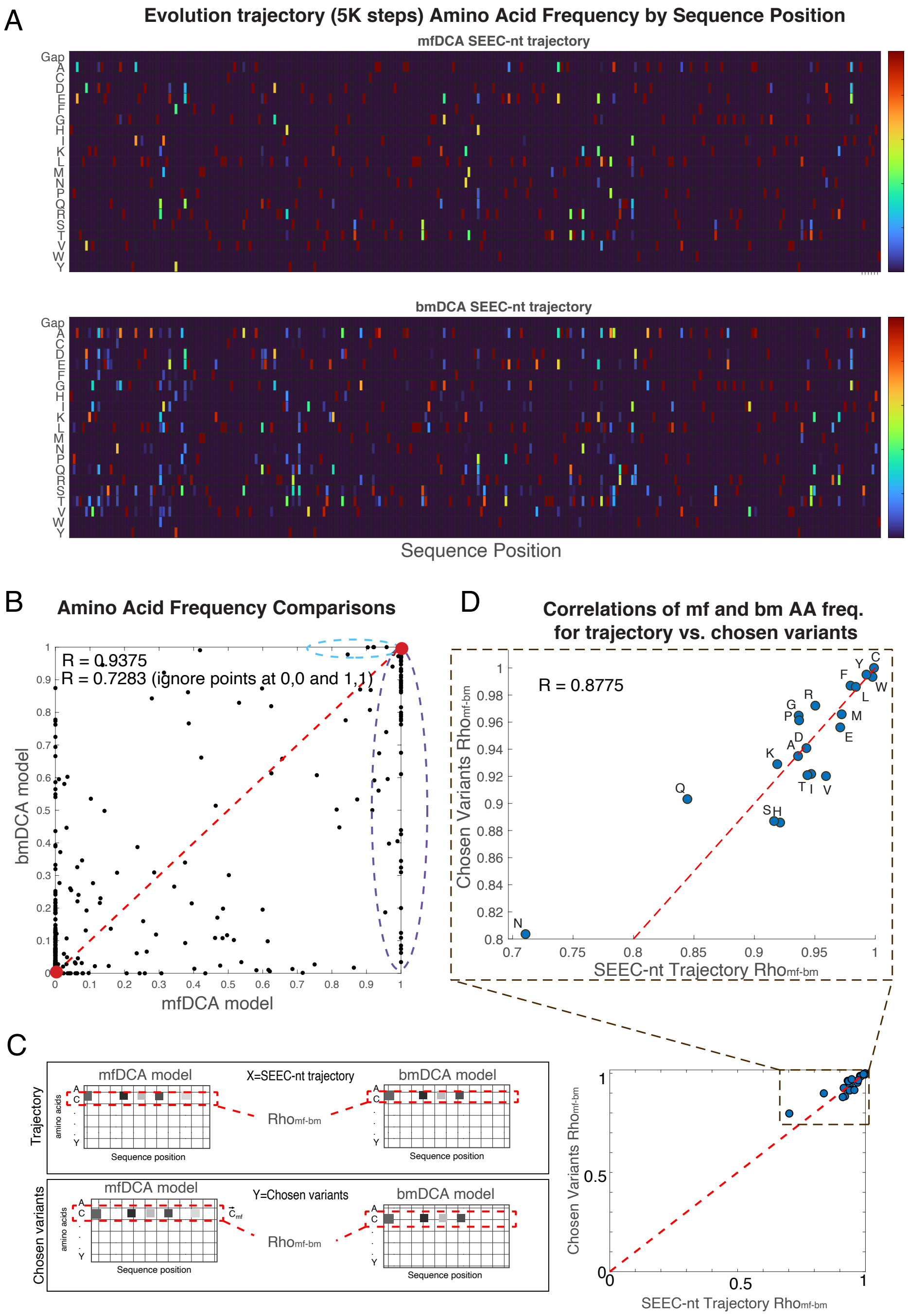


**Supplementary Figure 15:** Analysis of position-wise amino acid frequencies for the SEEC-nt evolutionary trajectories. (A) Heat maps of amino acid frequencies for each position in the sequence. Frequencies calculated across 5K variants generated during the simulation. Parameters used for the simulation came from either mfDCA (top) or bmDCA (bottom). (B) Comparing each point in the heat maps in a scatter plot to highlight the increased substitution rates of bmDCA over mfDCA. Purple oval features 57 positions that are conserved in the mfDCA-based trajectory but mutated in the bmDCA trajectory. In the light blue oval are 4 points representing positions that are conserved in bmDCA-based trajectory but mutated in the mfDCA trajectory. Red dots at 0,0 and 1,1 represent 5164 total points. There are 158 points at 1,1 and 5006 at 0,0. R values are correlation coefficients considering all points or the 359 points that remain after ignoring points at 1,1 and 0,0. (C) Overview of how each amino acid was individually analyzed for consistency of substitution frequency across both models and comparing the entire SEEC-nt trajectory to the subset of 26 variants chosen for experimental testing. (D) Inset and full scatter plot of correlation coefficients for each amino acid's frequencies across the sequence. Points are labeled with single letter amino acid code. Red dotted line of unity shown to depict minimal bias between full trajectory and chosen subset. Values are minimally biased

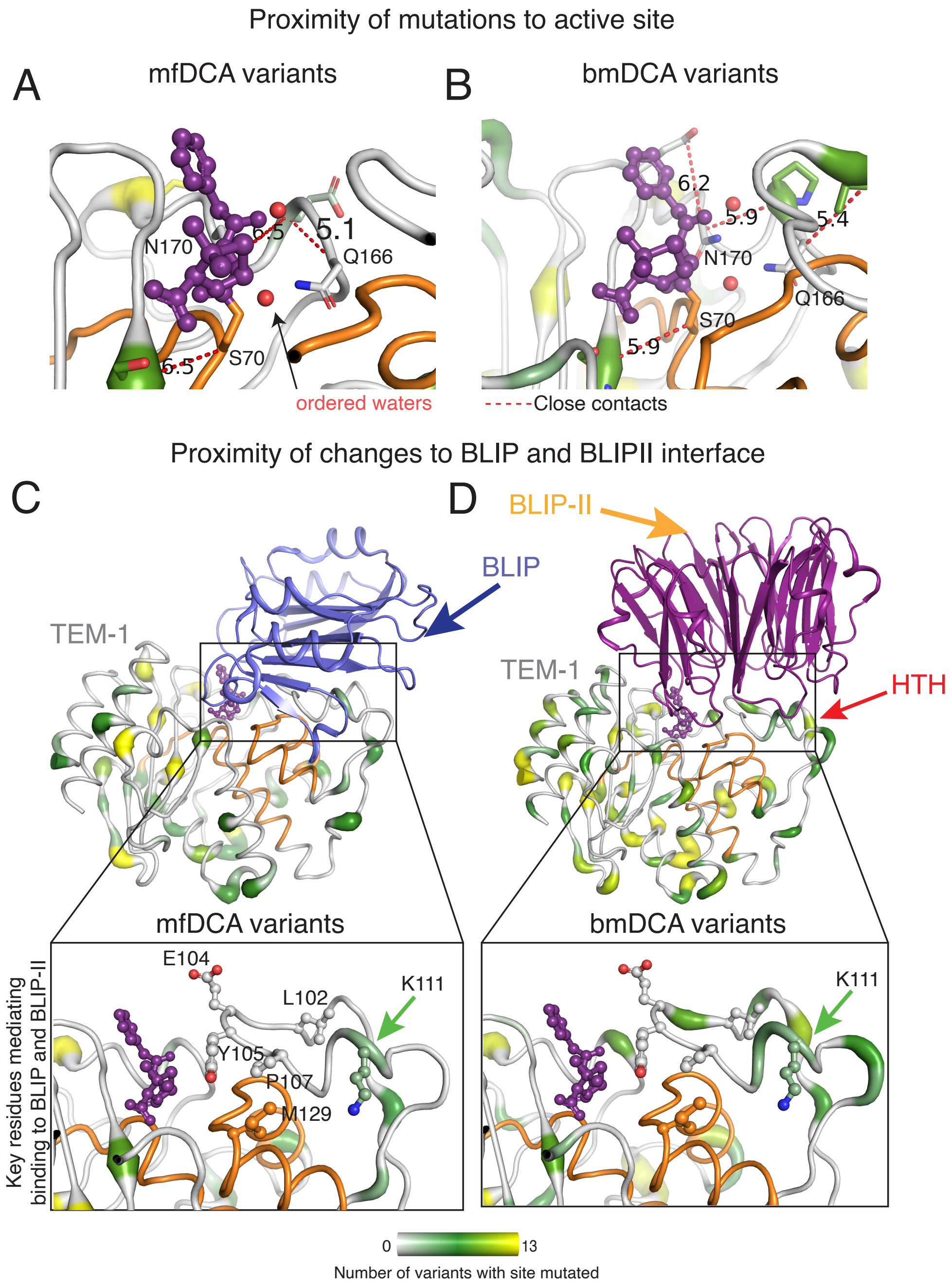


**Supplementary Figure 16:** Proximity of mutated sites to active site residues, S70, Q166 and N170. For the mfDCA variants (A), three mutated positions were identified as less than 8Åand are therefore contacts and for the bmDCA variants (B), there are four. None of these contacts are making direct chemical interactions and all of the substitutions are chemically similar except Pro167, which is mutated to Thr and Ala. Threonine is a conservative substitution, as it can back bond to Glu168 to help stabilize the kink that properly positions Q166 and N170 for catalysis. Structure depicted is PDBID 1FQG.(C and D) Complexes of E. coli TEM-1 with BLIP shown in blue (PDBID 3GMW, panel C) or BLIP-II shown in bright orange (D) As no structure of an E. coli TEM1-BLIP-II complex was found, the Bacillus anthracis beta-lactamase - Streptomyces exfoliatus BLIP-II complex structure (PDBID 3QHY) from was superimposed on the *E. coli* TEM-1 (1fqg) so that the location of the BLIP-II protein could be approximated. Despite significant sequence differences, the two beta-lactamases superimpose with 0.927ÅRMSD. Insets highlight specific residues that are key to mediating binding of BLIP and BLIP-II [2]. Lys111 is the only such key position that is mutated among the mf and bmDCA variants. In both cases it is mutated to Gln (see Fig. 5*A*,*B*). Orange regions in all panels highlight stretches of the structure where no mutations were found among the tested variants.

**Supplementary Table 1**
Sequences of Phase I variants generated from SEEC-AA using mfDCA and bmDCA inferred coupling and local field parameters.
**Supplementary Table 2**
Sequences of Phase II variants generated from SEEC-NT using mfDCA and bmDCA inferred coupling and local field parameters.
**Supplementary Table 3**
Phase II variants generated from SEEC-NT using mfDCA and bmDCA models with comparable percent identities.

### References


[1] Robert Fraczkiewicz and Werner Braun. Exact and efficient analytical calculation of the accessible surface areas and their gradients for macromolecules. *Journal of computational chemistry*, 19(3):319–333, 1998.

[2] Bartlomiej G Fryszczyn, Carolyn J Adamski, Nicholas G Brown, Kacie Rice, Wanzhi Huang, and Timothy Palzkill. Role of $\beta$-lactamase residues in a common interface for binding the structurally unrelated inhibitory proteins blip and blip-ii. *Protein Science*, 23(9):1235–1246, 2014.
