## Supplementary table 1 for "*In vivo* functional phenotypes from a computational epistatic model of evolution"

**Supplementary Table 1. SEEC-amino acid variants**

| Variant name | % ID to WT TEM1 | # of changes | Hamiltonian score/Temperature* | AA Sequence |
|---|---|---|---|---|
| Beg_mf_AA | 98.86 | 3 | -2042.20 | HPETLVKVKDAEDQLGARVGYIELDLNSGKILESFRPEERFPMMSTFKVLLCGAVLSRVDAGQEQLGRRIHYSQNDLVEYSPVTEKHLTDGMTVRELCSAAITMSDNTAANLLLTTIGGPKELTAFLHNTGDHVTRLDRWEPELNEAIPNDERDTTMPAAMATTLRKLLTGNLLTLASRQQLRDWMEADKVAGPLLRSALPAGWFIADKSGAGERGSRGIIAALGPDGKPSRIVVIYTTGSQATMDERNRQIAEIGASLIKHW |
| Mid_mf_AA | 70.72 | 77 | -2246.35 | HPETLVKVKDAEDQLGARVGYIELDLNSGKRWEEHRADERFPMCSTFKALACGAVLARVDRGEERLDRRVIGADDLVTYSPVTEKHVGVGMTLAELCEAAITLSDNTAGNLLLAAIGGPAGLTAFLRALGDDVTRLDRNEPELNEATPGDPRDTTTPAAMLATLRALLLGDALSAASRQQLRAWLVANKTGDALLRAGLPKGWRVGDKTGAGGAGTRNIVAVIWPPRRPSRIVVIYTTGSQATMDERNRQIAEIGASLIKHW |
| Late_mf_AA | 67.68 | 85 | -2277.43 | HPETLVKVKDAEDQLGARVGYIELDLNSGKRWEGHRADERFPMCSTFKFLAAAAVLARVDRGEERLDRRVVFGADDLVPYSPVTEKHVGVGMTLAELCEAAVTLSDNTAGNLLLASIGGPAGLTAFARSLGDPVTRLDRNEPELNEALPGDPRDTTTPAAMLEDLRALLLGDALSAASRQQLRAWLVANKTGDARLRAGLPKGWRVGDKTGTGGAGTTNDIAVIWPGRRPSRIVVIYTTGSQATMDERNRQIAEIGASLIKHW |
| Beg_bm_1_AA | 99.24 | 2 | -543.92 | HPETLVKVKDAEDQLGARVGYIELDLNSGKILESFRPEERFPMMSTFKVLLCGAVLSRVDAGQEQLGRRIHYSQNDLVEYSPVTEKHLTDGMTVRELCSAAITMSDNTAANLLLTTIGGPMELTAFLHNMGDHVTRLDRWEPELNEAIPNDERDTTMPAAMATTLRKLLTGELLTLASRQQLIDWMEADKVAGPLLRSALPAGWFIADKSGAGERGSRGDIAALGPDGKPSRIVVIYTTGSQATMDERNRQIAEIGASLIKHW |
| Beg_bm_2_AA | 98.86 | 3 | -543.37 | HPETLVKVKDAEDQLGARVGYIELDLNSGKILESFREEERFPMMSTFKVLLCGAVLSRVDAGQEQLGRRIHYSQNDLVEYSPVTEKHLTDGMTVRELCSAAITMSDNTAANLLLTTIGGPMELTAFLHNMGDHVTRLDRWEPELNEAIPNDERDTTMPAAMATTLRKLLTGELLTLASRQQLIDWMEADKVAGPLLRSALPAGWFIADKSGAGERGSRGDIAALGPDGKPSRIVVIYTTGSQATMDERNRQIAEIGASLIKHW |
| Beg_bm_3_AA | 98.48 | 4 | -545.88 | HPETLVKVKDAEDQLGARVGYIELDLNSGKILESFREEERFPMMSTFKVLLCGAVLSRVDAGQEQLGRRIHYSQNDLVEYSPVTEKHLTDGMTVRELCSAAITMSDNTAANLLLTTIGGPMELTAFLHNMGDHVTRLDRWEPELNEAIPNDERDTTMPAAMATTLRKLLTGELLTPASRQQLIDWMEADKVAGPLLRSALPAGWFIADKSGAGERGSRGDIAALGPDGKPSRIVVIYTTGSQATMDERNRQIAEIGASLIKHW |
| Mid_bm_1_AA | 67.30 | 86 | -462.24 | HPETLVKVKDAEDQLGARVGYIELDTQFERYRSDERFPMCSTFKVLLCAAVLSLIDKGIEQLERPVHFDQHDIVFYSPVTEAHVNAIMNKEEICSAADTTSDNTAANLVLDSIGGPPALTRFLWSPGDPATRLDRKEPFWNEAIPGDKRDTTTPAAMRTTLRLLLVGQWLSDKSRQQLAEWMRANKVATKLSRKSMPAGWVIADKFGMGGRDTRSIIAKFWPPGPPILISVVIYTTGSQATMDERNRQIAEIGASLIKHW |
| Mid_bm_2_AA | 67.68 | 85 | -465.13 | HPETLVKVKDAEDQLGARVGYIELDTQFERYRSDERFPMCSTFKVLLCAAVLSLIDKGIEQLERPVHFDQHDIVFYSPVTEAHVNAIMNVEEICSAADTTSDNTAANLVLDSIGGPPALTRFLWSPGDPATRLDRKEPFWNEAIPGDKRDTTTPAAMRTTLRLLLVGQWLSDKSRQQLAEWMRANKVATKLSRKSMPAGWVIADKFGMGGRDTRSIIAKFWPPGPPILISVVIYTTGSQATMDERNRQIAEIGASLIKHW |
| Mid_bm_3_AA | 67.30 | 86 | -465.51 | HPETLVKVKDAEDQLGARVGYIELDTQFERYRSDERFPMCSTFKVLLCAAVLSLIDKGIEQLERPVHFDAHDIVFYSPVTEAHVNAIMNVEEICSAADTTSDNTAANLVLDSIGGPPALTRFLWSPGDPATRLDRKEPFWNEAIPGDKRDTTTPAAMRTTLRLLLVGQWLSDKSRQQLAEWMRANKVATKLSRKSMPAGWVIADKFGMGGRDTRSIIAKFWPPGPPILISVVIYTTGSQATMDERNRQIAEIGASLIKHW |
| Late_well_bm_1_AA | 64.64 | 93 | -468.07 | HPETLVKVKDAEDQLGARVGYIELDTGRRWEGHRGDERFPMMSTFKSLLCGLVLSKQDSGSSDLDQRVSYCERELLEYSPVTEKRAPDGMTVSELCAATIDISDNTATNHLITSLGGHEALTDFARSIGDEKTRLDHYEPEGNEANNGDTRDTTTPAAMLATLKQLLLGDSLTLSSRQQLLMWMRANQVTMRTLRAAVPDDWAIADETGAGENGTRSIVGIDRPDGKPLIIAVVIYTTGSQATMDERNRQIAEIGASLIKHW |
| Late_well_bm_2_AA | 64.26 | 94 | -470.61 | HPETLVKVKDAEDQLGARVGYIELDTGRRWEGHRGDERFPMMSTFKSLLCGLVLSKQDSGSSDLDQRVSYCERELLEYSPVTEKRAPDGMTVSELCAATIDISDNTATNHLITSLGGHEALTDFARSIGDEKTRLDHYEPEGNEANNGDTRDTTTPAAMLATLKQLLLGDSLTPSSRQQLLMWMRANQVTMRTLRAAVPDDWAIADETGAGENGTRSIVGIDRPDGKPLIIAVVIYTTGSQATMDERNRQIAEIGASLIKHW |
| Late_well_bm_3_AA | 64.64 | 93 | -474.54 | HPETLVKVKDAEDQLGARVGYIELDTGRRWEGHRGDERFPMMSTFKSLLCGLVLSKVDSGSSDLDQRVSYCERELLEYSPVTEKRAPDGMTVSELCAATIDISDNTATNHLITSLGGHEALTDFARSIGDEKTRLDHYEPEGNEANNGDTRDTTTPAAMLATLKQLLLGDSLTPSSRQQLLMWMRANQVTMRTLRAAVPDDWAIADETGAGENGTRSIVGIDRPDGKPLIIAVVIYTTGSQATMDERNRQIAEIGASLIKHW |
| Late_ave_bm_1_AA | 63.88 | 95 | -451.79 | HPETLVKVKDAEDQLGARVGYIELDTGRRWEGHRCDDRFPMCSTFKSLLCGLILSRVDSGSSDLDERPSYCWRELVEYSPVTEKRAPDGMTMSELCAATIEISDNAATNLLIESMGGHQALTDFLRSIGDEKTRLDRYEPEGNEANNGDTRDTTTPAACAASLKQLLLGGSLLPSSRQRLLMWMRAARVTPRLLRAAVPDGWAIADETGAGENGTRGIWGIVRPDGKCLLIAVVIYTTGSQATMDERNRQIAEIGASLIKHW |
| Late_ave_bm_2_AA | 64.26 | 94 | -454.54 | HPETLVKVKDAEDQLGARVGYIELDTGRRWEGHRCDERFPMCSTFKSLLCGLILSRVDSGSSDLDERPSYCWRELVEYSPVTEKRAPDGMTMSELCAATIEISDNAATNLLIESMGGHQALTDFLRSIGDEKTRLDRYEPEGNEANNGDTRDTTTPAACAASLKQLLLGGSLLPSSRQRLLMWMRAARVTPRLLRAAVPDGWAIADETGAGENGTRGIWGIVRPDGKCLLIAVVIYTTGSQATMDERNRQIAEIGASLIKHW |
| Late_ave_bm_3_AA | 64.64 | 93 | -454.84 | HPETLVKVKDAEDQLGARVGYIELDTGRRWEGHRCDERFPMCSTFKSLLCGLILSRVDSGSSDLDERPSYCWRELVEYSPVTEKRAPDGMTMSELCAATIEISDNAATNLLIESMGGHQALTDFLRSIGDEKTRLDRYEPEGNEANNGDTRDTTTPAACAASLKQLLLGGSLLPSSRQRLLMWMRAARVTPRLLRAAVPDGWAIADETGAGENGSRGIWGIVRPDGKCLLIAVVIYTTGSQATMDERNRQIAEIGASLIKHW |
| Late_peak_bm_1_AA | 63.88 | 95 | -437.80 | HPETLVKVKDAEDQLGARVGYIELDTGKRWEGHRCDERFPMCSTFKSLLCGLILSRSDSGSSDLDERPRYCWRELVEYSPVAEKRAQDGMTMYELCAATIELSDNTATNLLIEHMGGHQGLTAFLRSIGDEWTLDRYEDEGNEANNGDTRDTTTPAACAHSLKQLALGGSLLPSSRQRLLMWMIAARVAPRLLRAAVPDGWAIADETGAGENGSRGIWGIVRPRGKPLLIAIVVIYTTGSQATMDERNRQIAEIGASLIKHW |
| Late_peak_bm_2_AA | 64.26 | 94 | -438.88 | HPETLVKVKDAEDQLGARVGYIELDTGKRWEGHRCDERFPMCSTFKSLLCGLILSRSDSGSSDLDERPRYCWRELVEYSPVAEKRAQDGMTMYELCAAAIELSDNTATNLLIEHMGGHQGLTAFLRSIGDEWTLDRYEDEGNEANNGDTRDTTTPAACAHSLKQLALGGSLLPSSRQRLLMWMIAARVAPRLLRAAVPDGWAIADETGAGENGSRGIWGIVRPRGKPLLIAIVVIYTTGSQATMDERNRQIAEIGASLIKHW |
| Late_peak_bm_3_AA | 63.88 | 95 | -437.00 | HPETLVKVKDAEDQLGARVGYIELDTGKRWEGHRCDERFPMCSTFKSLLCGLILSRSDSGSSDLDERPRYCWRELVEYSPVAEKRAQDGMTMYELCAAAIELSDNTATNLLIEHMGGHQGLTAFLRSIGDEWTLDRYEDEGNEANNGDTRDTSTPAACAHSLKQLALGGSLLPSSRQRLLMWMIAARVAPRLLRAAVPDGWAIADETGAGENGSRGIWGIVRPRGKPLLIAIVVIYTTGSQATMDERNRQIAEIGASLIKHW |

*mfDCA T = 1
*bmDCA T = 1
