## Supplementary table 3 for "*In vivo* functional phenotypes from a computational epistatic model of evolution"

**Supplementary Table 3.** **SEEC-nt equal percent identity variants across models**

| Model | Variant name | % ID to WT TEM1 | # of changes |
|---|---|---|---|
| bmDCA | Beg_bm_4_NT | 96.20 | 10 |
| mfDCA | Beg_mf_1_NT | 96.20 | 10 |
| | Beg_mf_2_NT | 96.58 | 9 |
| | Beg_mf_3_NT | 96.20 | 10 |
| bmDCA | Beg_bm_5_NT | 90.11 | 26 |
| | Beg_bm_6_NT | 90.11 | 26 |
| mfDCA | Mid_mf_1_NT | 90.87 | 24 |
| | Mid_mf_2_NT | 90.49 | 25 |
| | Mid_mf_3_NT | 90.11 | 26 |
| bmDCA | Beg_bm_3_NT | 86.31 | 36 |
| mfDCA | Late_mf_1_NT | 86.31 | 36 |
| | Late_mf_2_NT | 86.69 | 35 |
| | Late_mf_3_NT | 87.07 | 34 |
