## Supplementary table 2 for "*In vivo* functional phenotypes from a computational epistatic model of evolution"

**Supplementary Table 2.** **SEEC-nucleotide variants**

| Variant name | % ID to WT TEM1 | # of changes | Hamiltonian score/ | AA Sequence |
|---|---|---|---|---|
| Beg_mf_1_NT | 96.20 | 10 | -2316.64 | HPETLVKVKDAEDQLGARVGYIELDLNSGRILDSFRPEERFPMMSTFKVLLCGAVLSRVDAGQEQLGRRIHYSQNDLVEYSPVTEKHLTDGMTVRELCSAAITMSDNTAANLLLTTIGGPKELTAFLRNMGDHVTRLDRWEPELNEALPNDERDTTTPAAMATTLRKLLTGELLTLASRQQLIDWMEADKVAGPLLRSVLPAGWFIADKSGAGERGSRGIVAALGPDGKPSRIVVIYLTGTQATMDERNRQIAGIGASLIKHW |
| Beg_mf_2_NT | 96.58 | 9 | -2320.36 | HPETLVKVKDAEDQLGARVGYIELDLNSGRILDSFRPEERFPMMSTFKVLLCGAVLSRVDAGQEQLGRRIHYSQNDLVEYSPVTEKHLTDGMTVRELCSAAITMSDNTAANLLLTTIGGPKELTAFLRNMGDHVTRLDRWEPELNEALPNDERDTTTPAAMATTLRKLLTGELLTLASRQQLIDWMEADKVAGPLLRSVLPAGWFIADKSGAGERGSRGIVAALGPDGKPSRIVVIYLTGTQATMDERNRQIAEIGASLIKHW |
| Beg_mf_3_NT | 96.20 | 10 | -2317.61 | HPETLVKVKDAEDQLGARVGYIELDLNSGRILDSFRPEERFPMMSTFKVLLCGAVLSRVDAGQEQLGRRIHYSQNDLVEYSPVTEKHLTDGMTVRELCSAAITMSDNTAANLLLTTIGGPKELTAFLRNMGDHVTRLDRWEPELNEALPNDERDTTTPAAMATTLRRLLTGELLTLASRQQLIDWMEADKVAGPLLRSVLPAGWFIADKSGAGERGSRGIVAALGPDGKPSRIVVIYLTGTQATMDERNRQIAEIGASLIKHW |
| Beg_mf_4_NT | 97.34 | 7 | -2314.76 | HPETLVKVKDAEDQLGARVGYIELDLNSGRILESFRPEERFPMMSTFKVLLCGAVLSRVDAGQEQLGRRIHYSQNDLVEYSPVTEKHLTDGMTVRELCSAAITMSDNTAANLLLTTIGGPKELTAFLRNMGDHVTRLDRWEPELNEALPNDERDTTTPAAMATTLRKLLTGELLTLASRQQLIDWMEADKVAGPLLRSVLPAGWFIADKSGAGERGSRGIVAALGPDGKPSRIVVIYSTGSQATMDERNRQIAEIGASLIKHW |
| Beg_mf_5_NT | 95.82 | 11 | -2318.63 | HPETLVKVKDAEDQLGARVGYIELDLNSGRILDSYRPEERFPMMSTFKVLLCGAVLSRVDAGQEQLGRRIHYSQNDLVEYSPVTEKHLTDGMTVRELCSAAITMSDNTAANLLLTTIGGPKELTAFLRNMGDHVTRLDRWEPELNEALPNDERDTTTPAAMATTLRRLLTGELLTLASRQQLIDWMEADKVAGPLLRSVLPAGWFIADKSGAGERGSRGIVAALGPDGKPSRIVVIYLTGTQATMDERNRQIAEIGASLIKHW |
| Beg_mf_6_NT | 95.82 | 11 | -2325.72 | HPETLVKVKDAEDQLGARVGYIELDLNSGRILESYRPEERFPMMSTFKVLLCGAVLSRVDAGQEQLGRRIHYSQNDLVEYSPVTEKHLTDGMTVRELCSAAITMSDNTAANLLLTTIGGPKELTAFLRNMGDHVTRLDRWEPELNEALPNDERDTTTPAAMATTLRRLLTGELLTLASRQQLIDWMEADKVAGPLLRSVLPAGWFIADKTGAGERGSRGIVAALGPDGKPSRIVVIYLTGTQATMDERNRQIAEIGASLIKHW |
| Mid_mf_1_NT | 90.87 | 24 | -2325.56 | HPATLVKVKDAEDQLGARVGYIELDLNSGQILESYRPQERFPMMSTFKVLLCGAVLSRVDAGQEQLDRRIRYSQNDLVEYSPVTEKHLTDGMTVRELCSAAITMSDNTAANLLLSTIGGPKGLTAFLRKTGDHVTRLDRWEPELNEALPNDERDTTTPEAMATTLRTLLTGKLLTPASRQQLRDWMEADKVAGPLLRSVLPAGWFIADKTGAGERGSRGIVAALGPDGKPSRIVVIYLTGTQATMDERNRQIADIGASLIKHW |
| Mid_mf_2_NT | 90.49 | 25 | -2321.71 | HPATLVKVKDAEDQLGARVGYIELDLNSGQILESYRPQERFPMMSTFKVLLCGAVLSRVDAGQEQLDRRIRYSQNDLVEYSPVTEKHLTDGMTVRELCSAAITMSDNTAANLLLSTIGGPKGLTAFLRKTGDHVTRLDRWEPALNEALPNDERDTTTPEAMATTLRTLLTGKLLTPASRQQLRDWMEADKVAGPLLRSVLPAGWFIADKTGAGERGSRGIVAALGPDGKPSRIVVIYLTGTQATMDERNRQIADIGASLIKHW |
| Mid_mf_3_NT | 90.11 | 26 | -2321.42 | HPATLVKVKDAEDQLGARVGYIELDLNSGQILESYRPQERFPMMSTFKVLLCGAVLSRVDAGQEQLDRRIRYSQNDLVEYSPVTEKHLTDGMTVRELCSAAITMSDNTAANLLLSTIGGPKGLTAFLRKTGDHVTRLDRWEPALNEALPNDERDTTTPEAMARTLRTLLTGKLLTPASRQQLRDWMEADKVAGPLLRSVLPAGWFIADKTGAGERGSRGIVAALGPDGKPSRIVVIYLTGTQATMDERNRQIADIGASLIKHW |
| Mid_mf_4_NT | 92.40 | 20 | -2315.42 | HPETLVKVKDAEDQLGGRVGYIELDLNSGQILESYRPEERFPMMSTFKVLLCGAVLSRVDAGQEQLDRRIHYSQNDLVEYSPVTEKHLTDGMTVRELCSAAITMSDNTAANLLLSTIGGPKGLTAFLRNMGDHVTRLDRWEPELNEALPNDERDTTTPEAMARTLRKLLTGKLLTPASRQQLMDWMEADKVAGPLLRSVLPAGWFIADKTGAGERGSRGIVAALGPDDKPSRIVVIYLTGTQATMDERNRQIAEIGASLIKHW |
| Late_mf_1_NT | 86.31 | 36 | -2323.35 | HPATLDKVKDAEDQLGARVGYAELDLNSGQMLESYRPQERFPMMSTFKVLLCGAVLSRVDAGQEQLDRRIRYSQNDLVEYSPVTEQHLTDGMSVRELCSAAITMSDNTAANLLLSTIGGPQGLTAFLRKTGDQVTRLDRWEPELNEALPNDERDTTTPEAMARTLRTLLTGNVLTPTSQQQLQDWMEADKVAGPLLRSVLPAGWFIADKTGAGERGSRGIVAALGPDGKPSRIVVIYLTGTQATMDERNKQIAAIGASLIKHW |
| Late_mf_2_NT | 86.69 | 35 | -2324.81 | HPATLDKVKDAEDQLGARVGYAELDLNSGQMLESYRPQERFPMMSTFKVLLCGAVLSRVDAGQEQLDRRIRYSQNDLVEYSPVTEQHLTDGMSVRELCSAAITMSDNTAANLLLSTIGGPQGLTAFLRKTGDQVTRLDRWEPELNEALPNDERDTTTPEAMARTLRTLLTGNVLTPTSRQQLQDWMEADKVAGPLLRSVLPAGWFIADKTGAGERGSRGIVAALGPDGKPSRIVVIYLTGTQATMDERNKQIAAIGASLIKHW |
| Late_mf_3_NT | 87.07 | 34 | -2325.96 | HPATLDKVKDAEDQLGARVGYAELDLNSGQMLESYRPQERFPMMSTFKVLLCGAVLSRVDAGQEQLDRRIRYSQNDLVEYSPVTEQHLTDGMSVRELCSAAITMSDNTAANLLLSTIGGPQGLTAFLRKTGDQVTRLDRWEPELNEALPNDERDTTTPEAMARTLRTLLTGNVLTPASRQQLQDWMEADKVAGPLLRSVLPAGWFIADKTGAGERGSRGIVAALGPDGKPSRIVVIYLTGTQATMDERNKQIAAIGASLIKHW |
| Beg_bm_1_NT | 85.93 | 37 | -1187.58 | HPATLEKVKDAEEQLGARVGYSELNLSSGRILQSYRADERFPMMSTFKVLLCGAVLARVDAGQEQLGRRIHYTQNDLVEYSPVTEKHLTHGMTVRELCSAAITMSDNTAANLLLSKLGGPQGLTAFLRKTGDEVTRLDRWETELNEALPIDERDTTTPEAMAATLRKLLTGEALTLTSRQQLRDWMESDKVAGPLLRSALPAGWFIADKTGAGERGSRGIIAVLGPDGKPSRIVVIYMTGSQATMDERNRQIAEIGASLIKNW |
| Beg_bm_2_NT | 85.93 | 37 | -1189.28 | HPATLEKVKDAEEQLGARVGYSELNLSSGQILQSYRADERFPMMSTFKVLLCGAVLARVDAGQEQLGRRIHYTQNDLVEYSPVTEKHLTHGMTVRELCSAAITMSDNTAANLLLSKLGGPQGLTAFLRKTGDEVTRLDRWETELNEALPIDERDTTTPEAMAATLRKLLTGEALTLTSRQQLRDWMESDKVAGPLLRSALPAGWFIADKTGAGERGSRGIIAVLGPDGKPSRIVVIYMTGSQATMDERNRQIAEIGASLIKNW |
| Beg_bm_3_NT | 86.31 | 36 | -1193.97 | HPATLEKVKDAEEQLGARVGYSELDLSSGQILQSYRADERFPMMSTFKVLLCGAVLARVDAGQEQLGRRIHYTQNDLVEYSPVTEKHLTHGMTVRELCSAAITMSDNTAANLLLSKLGGPQGLTAFLRKTGDEVTRLDRWETELNEALPIDERDTTTPEAMAATLRKLLTGEALTLTSRQQLRDWMESDKVAGPLLRSALPAGWFIADKTGAGERGSRGIIAVLGPDGKPSRIVVIYMTGSQATMDERNRQIAEIGASLIKNW |
| Beg_bm_4_NT | 96.20 | 10 | -1189.82 | HPATLEKVKDAEDQLGARVGYSELDLNSGKILESFRPEERFPMMSTFKVLLCGAVLSRVDAGQEQLGRRIHYSQNDLIEYSPVTEKHLTDGMTVRELCSAAITMSDNTAANLLLTTIGGPKELTAFLHKTGDDVTRLDRWEPELNEAIPNDERDTTTPAAMATSLRKLLTGELLTLASRQQLIDWMESDKVAGPLLRSALPAGWFIADKSGAGERGSRGIIAALGPDGKPSRIVVIYTTGSQATMDERNRQIAEIGASLIKHW |
| Beg_bm_5_NT | 90.11 | 26 | -1211.07 | HPATLEKVKAAEDQLGARVGYSELDLNSGKVLESYRPEERFPMMSTFKVLLCGAVLARVDAGQEQLGRRIHYSQNDLIEYSPVTEKHLTDGMTVRELCSAAITMSDNTAANLLLTSIGGPQALTAFLRKTGDDVTRLDRWETELNEALPNDERDTTTPAAMASTLRNLLTGERLTLASRQQLIDWMESDKVAGPLLRSALPAGWFIADKSGAGERGSRGIIAALGPDGKPSRIVVIYMTGSQATMDERNKQIAGIGASLIKNW |
| Beg_bm_6_NT | 90.11 | 26 | -1212.26 | HPATLEKVKAAEDQLGGRVGYSELDLSSGKILESYRPEERFPMMSTFKVLLCGAVLARVDAGQEQLGRRIHYSQNDLVEYSPVTEKHLTHGMTVRELCSAAITMSDNTAANLLLTTIGGPQGLTAFLRKTGDDVTRLDRWETELNEALPNDERDTTTPAAMAATLRKLLTGERLTLASQQQLIDWMESDKVAGPLLRSALPAGWFIADKTGAGERGSRGIIAALGPDGKPSRIVVIYMTGSQATMDERNRQIAGIGASLIKNW |
| Mid_bm_1_NT | 82.51 | 46 | -1252.89 | HPATLDKGKFAESQLNARVGYAELDLASGWILESYRSNERFPMMSTFKVLLCGAVLSRVDAGQEQLDRRITYQQSDLIEYSPVSEKHLADGMTVGELCSAAITMSDNTAANLLLSTIGGPQGLTAFLRKTGDQVSRLDRWETELNEALPGDVRDTTTPEAMAQTLRNLLTGKALTATSRQQLIDWMEADKVAGPLLRSVLPAGWFIADKTGAGERGSRGIIAALGPDGKPSRIVVIYLTGSKATMDELNRQIAGIGASLIKHW |
| Mid_bm_2_NT | 82.13 | 47 | -1248.77 | HPATLDKGKFAESQLNARVGYAELDLASGWSLESYRSNERFPMMSTFKVLLCGAVLSRVDAGQEQLDRRITYQQSDLIEYSPVSEKHLADGMTVGELCSAAITMSDNTAANLLLSTIGGPQGLTAFLRKTGDQVSRLDRWETELNEALPGDVRDTTTPEAMAQTLRNLLTGKALTATSRQQLIDWMEADKVAGPLLRSVLPAGWFIADKTGAGERGSRGIIAALGPDGKPSRIVVIYLTGSKATMDELNRQIAGIGASLIKHW |
| Mid_bm_3_NT | 82.13 | 47 | -1242.13 | HPATLDKGKFAESQLNARVGYAELDLASGWSLESYRSNERFPMMSTFKVLLCGAVLSRVDAGQEQLDRRITYQQSDLIEYSPVSEKHLADGMTVGELCSAAITMSDNTAANLLLSTIGGPQGLTAFLRKTGDQVSRLDRWEAELNEALPGDVRDTTTPEAMAQTLRNLLTGKALTATSRQQLIDWMEADKVAGPLLRSVLPAGWFIADKTGAGERGSRGIIAALGPDGKPSRIVVIYLTGSKATMDELNRQIAGIGASLIKHW |
| Mid_bm_4_NT | 82.13 | 47 | -1254.15 | HPATLDKIKTSESQLNARVGYAELDLASGLILGSYRSDERFPMMSTFKVLLCGAVLSRVDAGQEQLDRRIAYRQSDLVEYSPVTKKHLADGMTVGELCSAAITMSDNTAANLLLSTIGGPQGLTAFLRKTGDPVTRLDRWEPELNEALPGDERDTTTPVAMAQTLRNLLTGKTLTAASEQQLLQWMEADKVAGPLLRSVLPAGWFIADKTGAGERGSRGIVAALGPDGKPSRIVVIYLTGSKATMDERNQQIADIGASLIKHW |
| Late_bm_1_NT | 78.71 | 56 | -1230.06 | HPATLDQIKQAESQLNGRVGYAELDLASGGILISYRPDERFPMMSTFKVLLCGAVLARVDAGLEQLDRRITYRQSDLIEYSPVTKQHLDDGMTVGELCSAAITMSDNTAANLLLSTIGGPQGLTAFLRKTGDRVSRLDRWEPELNEALPGDVRDTTTPEAMARTLRALLTGKTLTPTSKQQLIQWMEGDKVTGPLIRSVLPAGWFIADKSGAGERGSRGIVAALGPDGKPSRIVVIYLTGTSATMDERNEQIAGIGASLIKHW |
| Late_bm_2_NT | 78.33 | 57 | -1229.25 | HPATLDQIKQAESQLNGRVGYAELDLASGGILISYRLDERFPMMSTFKVLLCGAVLARVDAGLEQLDRRITYRQSDLIEYSPVTKQHLDDGMTVGELCSAAITMSDNTAANLLLSTIGGPQGLTAFLRKTGDRVSRLDRWEPELNEALPGDVRDTTTPEAMARTLRALLTGKTLTPTSKQQLIQWMEGDKVTGPLIRSVLPAGWFIADKSGAGERGSRGIVAALGPDGKPSRIVVIYLTGTSATMDERNEQIAGIGASLIKHW |
| Late_bm_3_NT | 77.95 | 58 | -1224.34 | HPATLDQIKQAESQLNGRVGYAELDLASGGILISYRLDERFPMMSTFKVLLCGAVLARVDAGLEQLDRRITYRQSDLIEYSPVTKQHLDDGMTVGELCSAAITMSDNTAANLLLSTIGGPQGLTAFLRKTGDRVSRLDRWEPELNEALPGDVRDTTTPEAMARTLRALLTGKTLTPTSKQQLIQWMEGDKVTGPLIRSVLPAGWFIADKSGAGDRGSRGIVAALGPDGKPSRIVVIYLTGTSATMDERNEQIAGIGASLIKHW |

*mfDCAT = 1.5
*bmDCAT = 0.75
